# Fat fish stay cool: Stress recovery and behavioral flexibility vary with nutritional state and predator exposure in the cichlid *Neolamprologus pulcher*

**DOI:** 10.1101/2025.04.02.646863

**Authors:** Stefan Fischer, Katharina Hirschenhauser, Barbara Taborsky, Leonida Fusani, Virginie Canoine, Sabine Tebbich

## Abstract

Behavioral flexibility plays an important role in adaptation to changing environments within an individual*’*s lifetime. Stress resilience influences flexibility, as short-term stress can enhance attention and memory, while prolonged stress may impair cognitive function. Nutritional state also plays a role, with overweight individuals often suffering greater consequences of chronic stress. The allostatic load model predicts that individuals under prolonged stress remain in a chronically activated state, potentially reducing flexibility.

Using a cichlid, we manipulated body condition and exposed individuals to occasional or frequent predator presentations. We then assessed stress responses and behavioral flexibility via a reversal learning test. Contrary to predictions, fish in a high nutritional state - but not overweight - that were frequently exposed to predators recovered rapidly from stress and exhibited enhanced flexibility. Our results provide direct evidence that nutritional state interacts with behavioral flexibility, revealing the costs of coping with repeated stressors.

We highlight a link between brief stress responses, characterized by rapid cortisol recovery, and behavioral flexibility. This association only emerged in well-nourished individuals facing frequent stressors, suggesting energy reserves buffer the costs of repeated stress. Thus, resilience benefits behavioral flexibility, while it is energetically costly, as a reduced diet hindered adaptation to environmental change.

## Introduction

Animals have to constantly cope with and adapt to stressors in the environment such as the presence of predators or changing environmental conditions (MacLeod et al., 2023). Stress resilience, i.e. the rapidity, efficiency, and completeness of the recovery of the stress response after a perturbation (sensu Wingfield, 2013) is a major component of the physiological adaptations for coping with environmental challenges. The hypothalamic-pituitary-adrenal/interrenal (HPA/HPI) axis plays a central role in mediating physiological stress resilience (Creel et al., 2013). This endocrine system controls the release of glucocorticoids (GCs), such as cortisol or corticosterone, which help mobilizing energy to cope with the stressor (Sapolsky et al., 2000). Stressors disrupt the homeostasis of an organism, and the concept of allostasis refers to the process by which the body reestablishes homeostasis through physiological or behavioral changes (McEwen, 2005; McEwen and Wingfield, 2003, 2010). Allostasis is costly and organisms accumulate allostatic load when exposed to stressors (Romero et al., 2009; Wright et al., 2023). An effective termination of the stress response is therefore essential to prevent the detrimental effects of prolonged exposure to stress hormones such as GCs (McEwen, 2005). Thus, acute stress responses are characterized by a short activation of the HPA/HPI axis with the result of increased GC secretion from the adrenals, followed by a swift decrease and return to normal conditions. If the stressors occur repeatedly, but only occasionally, there is sufficient time for the organism to quickly restore homeostasis, thus keeping the cost of allostasis low. An acute stress response is beneficial, because it effectively activates physiological and behavioral processes to cope with the stressor. Such acute stress responses are adaptive and important for survival with positive effects on health and cognition (de Kloet et al., 2005; Sandi and Pinelo-Nava, 2007; Schwabe et al., 2012).

In contrast, prolonged stress responses are characterized by a long activation of the HPA/HPI axis with persistently elevated GC concentrations (Dickens and Romero, 2013). This persistent accumulation of allostatic load is energetically costly and may lead to *‘*wear and tear*’* of the organism*’*s body (Romero et al., 2009). If stressors occur frequently and the organisms cannot recover, these wear and tear costs accumulate and may contribute to a range of health problems and impair cognition (D*’*Amico et al., 2020).

Behavioral flexibility is the capacity to adapt behavior in response to changing environmental conditions (see Burkart et al., 2017; Diamond, 2013; Sol et al., 2002). Arguably, cognition is among the most important determinants of behavioral flexibility because learned behavior can be reversed, allowing for significant adaptability (Tebbich et al., 2010). While exposure to acute stressors may enhance behavioral flexibility (Bryce and Howland, 2015; Graybeal et al., 2011; Thai et al., 2013), prolonged or frequent stressors impair behavioral flexibility, possibly as a result of persistently elevated stress responses (George et al., 2015).

The type and the duration of the stressor, as well as the energetic reserves of the organism determine whether a stressor elicits an acute or prolonged stress response (Canoine et al., 2002; McEwen, 2005; McEwen and Wingfield, 2003). Previous studies proposed that overnutrition results in the accumulation of excessive baseline energy reserves that fuel prolonged allostasis; therefore the assumption was that obese organisms would be more susceptible to the effects of prolonged stress responses than lean organisms (McEwen, 2005; McEwen and Wingfield, 2003). The consequences can be severe. For instance, in humans chronic stress has been associated to a heightened risk for obesity, which in turn results in negative health consequences such as depression and an increased risk of inflammations (Benson et al., 2009; Kühnel et al., 2023; Petry et al., 2008).

A resilient organism can quickly overcome the allostatic load induced by a stressor and returns to homeostasis. Resilience is affected by several factors including the energetic reserves and life-history characteristics of the species (Wingfield, 2013). In humans, resilient stress response patterns reveal important health considerations, e. g. poor stress recovery correlates with poor cardiovascular status (Chida and Steptoe, 2010) and obesity has been linked to lower stress responsiveness and impaired cognition (Jones et al., 2012; Mujica-Parodi et al., 2009). Characterizing the specific factors that determine resilience in humans is a medically important question. However, identifying these important and diverse factors is extremely difficult, particularly in humans, due to the limits of experimental manipulations (Chbeir and Carrión, 2023). Thus, more research about the interaction between nutrition, stress responses and cognition is needed, particularly in animal models. So far, studies in laboratory rodents and birds showed rather mixed results (Kitaysky et al., 1999; Lynn et al., 2003; Schultner et al., 2013; Spencer and Tilbrook, 2009) with positive, negative, or no relationships between body condition and glucocorticoid responsiveness. We are not aware of any previous studies that have experimentally investigated the influence of nutritional state on the recovery pattern of stress responses and the resulting consequences for behavioral flexibility.

Fish and in particular cichlids are excellent models to further develop the research fields of cognition and stress. The physiological mechanisms to cope with stressors are evolutionary conserved among vertebrates (Nyman et al., 2018) and cichlids display a broad repertoire of socio-cognitive abilities (Jordan et al., 2021) showing a high degree of behavioral flexibility (Fischer et al., 2021; Reyes-Contreras and Taborsky, 2022). Standard procedures for experimental stress exposure are available, and individuals can be easily tagged to obtain individual characteristics (e.g. body condition) (Jungwirth et al., 2019; Watve et al., 2019). Combined with detailed physiological measurements, this allows us to use both functional and mechanistic approaches in experiments with cichlids.

*Neolamprologus pulcher* is a highly social cichlid, in which numerous previous studies have demonstrated the significance of the stress axis and stress reactivity for its social behavior and cognition (Antunes et al., 2021; Nyman et al., 2017, 2018; Reyes-Contreras et al., 2019; Reyes-Contreras and Taborsky, 2022; Taborsky et al., 2013). Thus, these fish are a well-suited model to test how the frequency of stressors and nutritional status influence stress response patterns and behavioral flexibility.

Here we performed an experiment in *N. pulcher* to test the hypothesis that the frequency of stressors and the nutritional state change individual cortisol response patterns which ultimately influences behavioral flexibility. To test this hypothesis, we manipulated the nutritional status of individuals while exposing them to different frequencies of recurrent stressors in the form of occasional or frequent predator presentations, assuming that the latter represents a prolonged stressor. At the end of the experiment, we assessed individual stress response patterns to a novel stressor and behavioral flexibility using a reversal learning test paradigm (Fischer et al., 2021). Based on the allostatic load model we had two specific predictions: (1) Frequent exposure to stressors and overnutrition separately induce prolonged allostatic states including persistently prolonged high cortisol concentrations and impaired behavioral flexibility, whereas an occasional exposure to stressors and a lower nutritional status induce brief allostatic states. (2) When these two factors are combined, overweight individuals are particularly susceptible to the negative effects of frequent exposure to stress, compared to lean individuals.

## Methods

### Subjects and housing

The experiment was conducted at the Konrad Lorenz Institute of Ethology, University of Veterinary Medicine, Vienna, Austria, under the license (ETK-101/06/2019). All procedures adhered to the highest standard for the treatment of animals and follow the guidelines for the ethical treatment of nonhuman animals in behavioral research and teaching (ASAB Ethical Committee, 2023). An ethical protocol was applied (see below) whenever signs of aggression were observed to avoid injuries or fatal outcomes. During the experiment one fish died which was caused by a precondition and not due to stress during any of the procedures. All experimental tanks were equipped with a 2 cm layer of sand, a biological filter, and a heater. The water temperature was kept at 27 ± 1°C with a light-dark regime of 13:11 h and a dimmed-light phase of 10 min. All biochemical parameters were close to the water conditions at southern Lake Tanganyika.

The study species was the cooperatively breeding *N. pulcher*, which is endemic to Lake Tanganyika. It lives in stable social groups always including a dominant breeding pair and related or unrelated helpers (Balshine et al., 2001; Heg et al., 2005; Taborsky, 1984, 1985). As an ecologically relevant stress stimulus, we used the piscivorous cichlid *Lepidiolamprologus elongatus*, which coexists with *N. pulcher* (Ochi and Yanagisawa, 1998). *L. elongatus* preys on *N. pulcher* of all sizes except for the eggs and is one of its most significant predators (Groenewoud et al., 2016).

### Experimental design

For this experiment we used 126 (63 male and 63 female) adult *N. pulcher* randomly selected from the institutès stock population. Before the experiment all fish were of similar size and body mass (size: 6.94 ± 0.03; body mass: 10.21 ± 0.14 g [mean ± SE]). The experiment was run in three separate blocks with 42 (21 females and 21 males) fish each that were housed in sex separated groups of 3-7 individuals (see below and Supplementary Fig. S1). A block design with three blocks was chosen for logistical reasons and we controlled potential temporal effects by always including all six treatments in each experimental block. To test the hypothesis outlined in the introduction, 30 groups of fish were created in total and each group received one of the following six treatments, resulting in five groups and a total of 21 test fish per treatment (Fig. 1 and Supplementary Fig. S2): (1) increased diet, no predators (IN); (2) increased diet, occasional predators [IO]; (3) increased diet, frequent predators [IF]; (4) reduced diet, no predators [RN]; (5) reduced diet, occasional predators [RO]; and (6) reduced diet, frequent predators [RF]. At the end of the treatments, we randomly selected 96 test fish from each group and assessed either their stress response patterns to a novel stressor (1 or 2 per group, Fig. S1) or their behavioural flexibility (1, 2 or 4 per group, Fig. S1). From the remaining fish (N=30, 1 per group) we collected the entire brains for a separate study that will be presented elsewhere. The experiment consisted of four phases: (1) the feeding regime, (2) predator presentations, (3) reversal learning task, and (4) hormone sampling. Each block of four phases lasted nine weeks (Fig. 1). Throughout the experiment we constantly monitored signs of aggression between group members. In cases where we observed that fish had received overt aggression (e.g. they had fins with frayed edges, were swimming close to the water surface or in the corner of the tank), we immediately placed a transparent divider with holes in the tank for the remainder of the experiment to physically separate fish from other group members without interfering with visual or olfactory contact (for more details see Supplementary Methods S1).

**Fig. 1:**
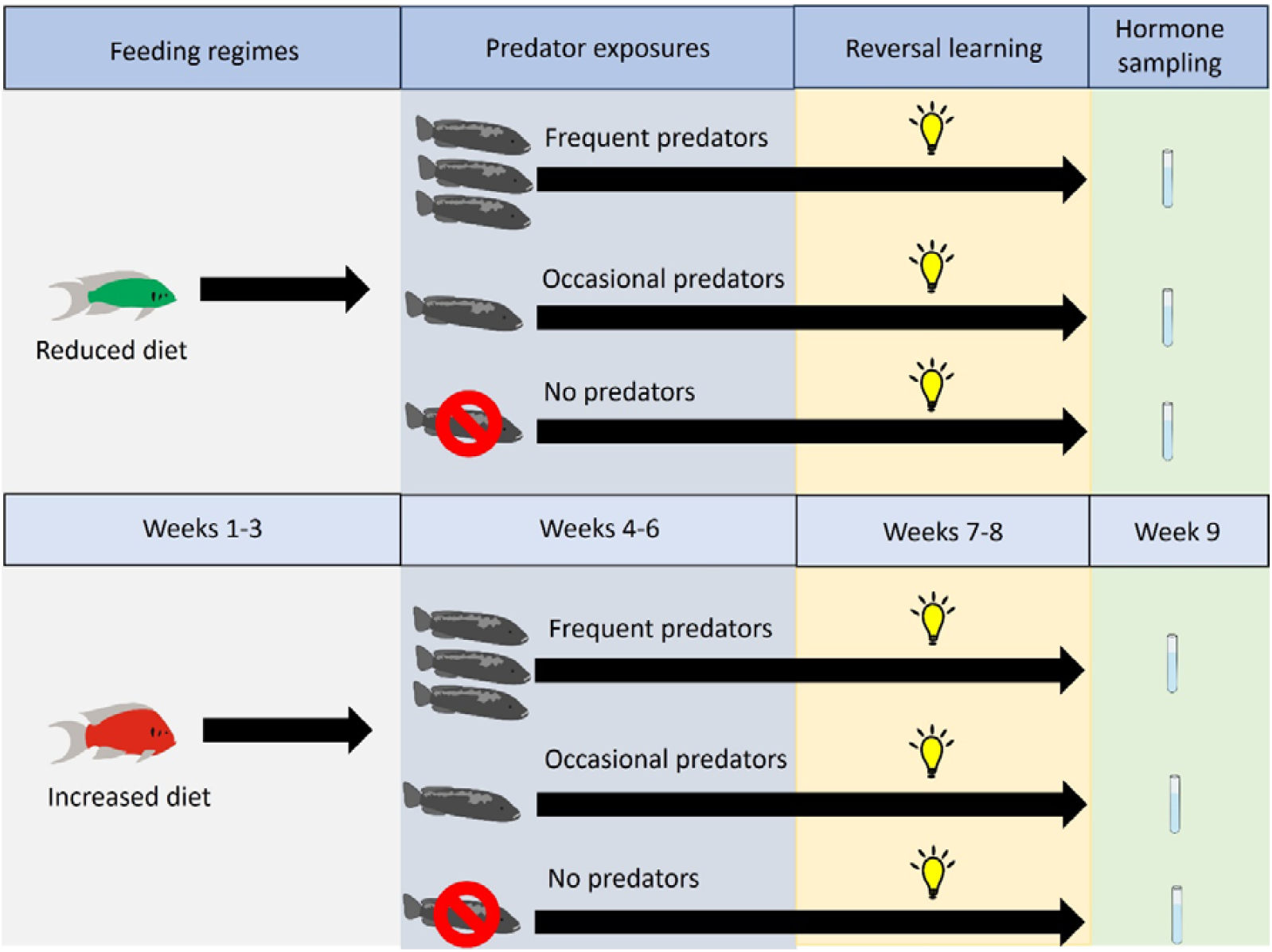
Timeline of the experiment. The experiment was done in three blocks (not shown) and each block lasted nine weeks which were separated into four phases. In the first phase, fish were kept either on an increased or reduced diet for three weeks. In the second phase, while continuing the feeding regimes, fish were exposed to three different predator exposures: (1) a control exposure without any predators, (2) a occasional or (3) a frequent exposure. In the third phase, we assessed behavioural flexibility using a reversal learning task. In the fourth phase, we collected water samples after exposure to a new stressor to measure stress response patterns in terms of cortisol concentrations.

### Feeding regime

Each group was assigned to one of two feeding regimes: (1) an increased or (2) a reduced diet. Fish were fed with commercial JBL Stick M for cichlids® once a day six times per week. The increased and reduced diet consisted of either 1.5 or 0.5 sticks per fish respectively (see Supplementary Methods S2 for results of a pilot experiment which confirms that the two diets induced fast and long-lasting changes in body condition). We chose JBL Stick M for cichlids® as food because of two reasons: (1) Individual sticks within a package are uniform and have nearly the same weight (0.045 ± 0.002g [mean ± SEM], N=10), which enabled us to create the two different food regimes by varying the number of sticks provided to the groups. (2) The sticks float on the water surface for a prolonged time and this ensured that all group members received an equal amount of food.

In each block the experiment started with the assignment of each group to its respective feeding regime and fish were fed the diet until the start of the hormone sampling phase (phase 4). During feeding we always placed two transparent dividers inside the tanks to assure that all fish within the same group received similar amounts of food. The dividers were arranged in a way to split up larger groups of fish into individual fish or small groups of 2-3 individuals depending on the initial group size. Particular attention was paid to avoid the repeated formation of small feeding groups that consisted of the same individuals. Dividers were left in place until most of the food was consumed and removed thereafter.

### Predator presentations

After fish had received their respective dietary treatments for three weeks, each group was assigned to one of three predator presentations. In blocks 2 and 3, two groups were assigned to the same predator presentations resulting in the same number of fish per treatment overall. All predator presentations consisted of a 5 min video playback of either (1) a recording of an empty tank, without any predators visible, or (2) a recording of an attacking predator. To manipulate the frequency of predator exposure, we presented the recording of the attacking predator either three times per week (occasional exposure) or three times per day (frequent exposure).

We presented the video playbacks using a 25.7cm [10.1⍰⍰] display using a laptop (Lenovo Ideapad D330 81MD-0006 Tablet Top). Videos were pre-recorded and validated to elicit similar responses as live predators (Watve et al., 2019). When presenting a video of an attacking predator, we randomly selected one of the four available videos and randomly placed the screen either left or right next to the experimental tank. To ensure that fish perceive the predators also by olfactory cues, we transferred 20ml of predator water to create a more naturalistic experience of an approaching predator. Predator water was always derived from one of two tanks (400L) containing five adult *L. elongatus* each. To avoid that test fish habituated to the predator presentations we spread out the presentations semi-randomly throughout the entire day and extended it with additional standardized stress manipulations (see Supplementary methods S3). To control for potential disturbances caused by adding predator water during the presentation of predator recordings, we also added 20ml of water from tanks containing groups of *N. pulcher* not used in the experiment during the presentation of control recordings.

### Reversal learning task

We used a well-established reversal learning task to measure behavioral flexibility (Fischer et al., 2021; Reyes-Contreras and Taborsky, 2022). For this, fish first had to learn a visual discrimination task and subsequently perform the reverse of the previously learned reward contingency. From each treatment group, we randomly selected 10 fish to be tested individually (in total N=60 test fish [Block 1: N=22; Block 2,3: N=18]). The task was separated into three phases: (1) pre-training and habituation phase, (2) training and color discrimination phase and (3) reversal learning phase. One fish died before the start of the training phase reducing the overall sample size to N=59.

#### (1) Pre-training and habituation phase

During the last two weeks of the predator presentations, we conducted the pre-training phase to habituate the fish to the test apparatus. The test apparatus consisted of an experimental dark grey slate placed on the sandy bottom of the tank. The slate contained 16 wells arranged in 4 columns, which could be covered with colored discs of 15 mm diameter. During the normal feeding schedules (as described above), when the groups were separated into smaller feeding groups, we also placed an experimental slate in each separation. We hid a small amount of food (crushed JBL Stick M for cichlids, according to their dietary treatments) in some of the wells which were either uncovered or only partially covered with a colored disc (irrespective of the color). During the two-week period we consecutively increased the coverage of the wells by the colored discs (25%, 50%, 75%, 100%) (see Buechel et al., 2018).

#### (2) Training phase

After the two-week pre-training and habituation phase we randomly selected fish from each group for the reversal learning task. Each fish was weighed and measured as described above and transported to an experimental training tank (30 x 30 x 50 cm). Each training tank was equipped with a 2cm layer of sand, a biological filter, a heater, and a half clay flowerpot as shelter. Each training tank was divided into a home compartment (2/3 of the tank) and an experimental compartment (1/3 of the tank). The two compartments were separated by an opaque barrier attached to strings on pulleys, allowing the observer to lift the barrier remotely while hiding behind a curtain to minimize disturbance of the fish during the procedure. The training tanks were placed side by side to allow the fish to interact socially, ensuring the welfare of the experimental fish and encouraging their participation during the tests. During tests we placed opaque barriers between the training tanks which avoided that fish in neighboring tanks influenced or eavesdropped on the behavior of the focal fish. Each fish was allowed to acclimate overnight to the new environment before the training phase started. Human observers were hidden behind a curtain, and observations were made blind to the experimental treatments. All observations were recorded using a web cam placed on a tripod in front of the curtain which was connected via a USB cable to a laptop at the other side of the curtain. This set-up allowed us to record and directly observe each test without the need to lift the curtain and potentially disturb the fish.

As soon as the fish had habituated to the novel experimental tank, we started to train them to access food from the feeding slate. We placed a feeding slate in the experimental compartment, removed the barrier to the home compartment and allowed the fish to access the experimental compartment and the food on the feeding slate. In the first training trial, a small green and a blue plastic disc were placed again next to one of the holes containing a reward (a red mosquito larvae). On each successive trial, the discs were moved further until they entirely covered the holes with the rewards. The fish had to learn to remove the disc by pushing it away to reach the reward. We always performed six trials per day for a maximum of 4 days or until fish successfully removed a disc to reach a reward that was entirely covered. Once they successfully removed a disc (N=56) or repeatedly entered the experimental compartment to feed on the reward (N=2), fish proceeded to the acquisition phase on the next day. One fish was excluded from analysis because it didn*’*t enter the compartment to get the reward during habituation.

#### (3) Color discrimination in the acquisition phase

In this phase, fish (N=58) were given the choice between two colored discs (blue and green), one with an accessible reward and one with no reward underneath. To prevent fish from uncovering the non-rewarded hole, we attached a pin below the unrewarded disc, which made it hard for the test fish to push the disc away and uncover the well. We placed a few drops of water which had previously contained mosquito larvae but not the actual larvae, into the hole covered by the unrewarded disc. This ensured that fish used visual cues (i.e. the color of the discs) and not any other cues (i.e. the smell of the reward) to discriminate between the rewarded and unrewarded discs. The rewarded color was randomly assigned so that half of the fish always found the reward underneath the blue disc, while the other half of the fish found the reward underneath the green disc. We always placed one disc on the left and the other disc on the right side of the slate on a randomly chosen well. On the next trial, the disc previously placed on the right side was placed on the left side on a randomly selected well and vice versa, and we continued to switch sides on each subsequent trial.

After the feeding slate was prepared, the observer moved behind the curtain and lifted the barrier between the experimental and the home compartment using a string. The angle of the video camera was positioned in a way that allowed us to test 3 fish at the same time. After the barrier was lifted, each fish was allowed to explore the experimental compartment for a maximum of 3 minutes. In a successful trial, the fish first uncovered the rewarded disc and accessed the reward. Immediately afterwards the observer closed the barrier and the fish moved back into the home compartment waiting for the next trial to start. In an unsuccessful trial, the fish either touched the unrewarded disc or did not touch any discs until the end of the trial. If the fish first touched the unrewarded disc or did not touch any discs, the observer did not close the barrier allowing the fish more time to move the rewarded disc and to uncover the reward. If the fish did not move the correct disc within an additional 2-minutes, the observer closed the barrier and moved the fish back into the home compartment. Afterwards, the observer moved the rewarded disc to reveal the reward underneath and opened the barrier until the fish consumed the reward or for a maximum of 2 min. If the fish did not consume the reward within these additional 2 minutes, the observer closed the barrier again and the fish moved back into the home compartment. The uneaten reward was transferred from the feeding slate into the home compartment, and the fish could consume the reward while the preparations for the next trial took place. This ensured that all fish consumed the same amount of food during each testing day. This procedure was repeated for six consecutive trials per day until the fish reached the learning criterion. We combined each set of 12 consecutive trials to one *“*block*”* of trials. As we did 6 trials per day, a block lasted at least for two days. To reach the learning criterion, the fish had to first choose the rewarded disc in 10 out of 12 trials (i.e., 80% of correct choices per block) with a maximum of one unsuccessful choice per day. If fish did not reach the learning criterion at the end of a block of 12 trials, the fish entered the next block of 12 trials. This procedure was repeated until the fish either reached the learning criterion or had passed five blocks (60 trials) without reaching the criterion. Once a fish reached the learning criterion in the acquisition phase, we started the reversal phase on the next day. Two fish did not reach the learning criteria and did not proceed to the reversal phase, reducing the sample size to N=56 for the next phase.

#### (3) Reversal learning phase

In this phase we reversed the rewarded and unrewarded color discs, so that the previously unrewarded disc became the rewarded one and vice versa. Otherwise, all procedures and recordings were done as described for the acquisition phase. The reversal phase ended when a fish had reached the learning criterion or after five blocks. All fish tested in the reversal phase reached the learning criterion.

### Hormone sampling

After the reversal learning task, we selected 1-2 fish from each group (N=36 fish, Block 1: N=11; Block 2: N=13: Block 3: N=12, Supplementary Fig. S1) that had received the food and predator treatments but not the reversal learning task. At the same time all predator presentations stopped, and all groups were transferred to an intermediate food diet (1 stick of JBL Stick M for cichlids® per fish.) The apparatus (see below) enabled the collection of water samples from two fish at the same time. We extended the static holding water method (Scott et al., 2008) with a continuous water flow that allows the repeated water sampling without further disturbing the fish (Antunes et al., 2021). The collections for each pair of fish took place over four consecutive days which allowed us to quantify cortisol concentrations (as ng cortisol/L) immediately after a new stressor (i.e. catching and handling of the fish on day 1) as well as cortisol concentrations after fish were habituated to this stressor, i.e. the same catching and handling procedures on day 4 (see below). The involved procedures, including catching, measuring, weighing, and placing the fish in a novel container are perceived as stressful by the experimental animals (Antunes et al., 2021; Wong et al., 2008). However, another study showed that *N. pulcher* habituate to this procedure after three days, which allows the collection of cortisol concentrations of fish before and after habituation (Antunes et al., 2021) and thus serves, as a reliable baseline measure.

On the first hormone sampling day we caught two fish and measured their body sizes and body masses. We then placed each of the two fish individually in a sampling container filled with 500ml of water and collected four 500ml water samples over 2h. All water samples were collected at the same time in the afternoon (after 14:00h) to control for diurnal variation of cortisol (Antunes et al., 2021). On two consecutive days, we caught the same two fish and transferred them to the sampling containers filled with 500ml of water for 2h but did not collect any water samples. This ensured that fish habituated to the catching and handling procedures of the hormone sampling. On the fourth day, we repeated the same procedures, including measuring and weighing the fish and collected another set of four water samples for 2h (see Supplementary Fig. S3). In block 1, due to logistical constraints, we only collected 2 water samples per day, instead of 8, and we decided to always collect the 3^rd^ and 4^th^ water samples on the 1^st^ and 4^th^ day respectively. All other procedures were the same for all three blocks. This resulted in a total of 240 water samples.

### Hormone measurement

#### Apparatus

To repeatedly and minimally invasively sample individual cortisol levels, we used a modification of the holding-water method previously used and validated for *N. pulcher* (Antunes et al., 2021). Exact details of the apparatus can be found in Supplementary Methods S4. In short, we used a water flow-through system to collect four water samples per fish throughout a 2h sampling period.

#### Cortisol measures

Prior analyzing cortisol levels, an extraction was performed as following: Water samples were defrosted overnight and filtered in the morning of the next day (paper filter 1/2595 grade, diameter 320 mm, Whatman, Sigma-Aldrich, Switzerland) to remove solid particles, and lipophilic compounds were extracted using solid phase extraction (SPE). After the hormone samples were loaded onto a cartridge (SPE column Isolute C18 (EC), 500 mg/6mL, Biotage, Sweden), the cartridges were stored at - 20 ⍰C until further processing at the University of Vienna (see Supplementary Methods S5). Of the 36 sampled test fish, we were not able to measure cortisol from 2 fish in any of the collected water samples, reducing the overall sample size to 34 fish.

### Statistical analysis

For statistical analyses we used R 4.3.3 (R Core Team, 2024) and the packages lme4 (Bates et al., 2015), and afex (Singmann et al., 2021). We used linear mixed effects models (LMM) and generalized linear mixed effects models (GLMM). To obtain p-values, we used either the mixed() function in the package *‘*afex*’* with a Satterthwaite*’*s approximation for degrees of freedom, or the drop1() function to do a likelihood ratio test using an F statistics for LMM or an X^2^ statistics for GLMM. Interactions and covariates were stepwise removed if non-significant (Engqvist, 2005) and reported p-values refer to the final model without non-significant interactions. An overview of all full models is provided in Supplementary Table S3. To account for familiarity of fish kept in the same group tanks and for potential variations between blocks we always included group tank ID and block ID as random factors. Based on our predictions and the rationale outlined in the introduction, we aimed to investigate the influence of nutritional state and not feeding regime *per se* on stress reactivity and behavioral flexibility. To compare fish with high and low body condition, we classified them as *‘*high nutritional state*’* or *‘*low nutritional state*’* depending on whether their body condition was higher or lower than the mean body condition of all fish tested in this study (3.02 ± 0.03 [mean ± SE]). We verified nevertheless that the feeding regimes had the desired effect and found that fish on the increased diet gained more weight and grew more throughout the experiment than fish on the reduced diet (see Supplementary Table S4, Figs. S6, S7). Consequently, the majority (71%) of fish on the increased diet had above-average body condition and were classified as *‘*high nutritional state*’* whereas the remaining fish (29%) were classified as *‘*low nutritional state*’*. The majority (74%) of fish on the reduced diet had below average body condition and were classified as *‘*low nutritional state*’* whereas the remaining fish (26%) were classified as *‘*high nutritional state*’*.

To analyze stress response patterns, we first classified each individual as a brief or prolonged responder based on their cortisol pattern in response to the hormone sampling on the first day. Brief responders had shortly elevated cortisol peaks whereas prolonged responders had persistently elevated cortisol concentrations within the 2h sampling period. We used a GLMM with a binomial error distribution to analyze whether the probability to display a brief stress response is influenced by body condition or predator exposures.

To estimate the duration of individual stress response patterns, we calculated an index of stress recovery on the first day of sampling. This index was calculated as the difference between the individual peak cortisol concentration and the final value after 2h (at sampling point 4). Thus, a higher stress recovery index indicates a stress response pattern characterized by a short cortisol peak with a rapid decrease of cortisol within 2h. A lower recovery value indicates a stress response pattern characterized by prolonged periods of high cortisol levels with a slow decrease within 2h. A recovery value of zero indicates a stress response pattern characterized by prolonged periods of high cortisol levels with no decrease within 2h. We discovered an influence of predator exposure on the likelihood of displaying a brief or prolonged stress response (see *Results*). This result was clearly driven by fish with high body condition although body condition was not significant in this analysis (see *Results* and Fig. 2). To assess whether the duration of the stress response is determined by the body condition of the fish, we used a LMM with stress recovery as the dependent variable and the actual body condition as a covariate. We conducted this analysis after confirming that fish exposed to frequent predators were more likely to show a brief stress response pattern. Therefore, to increase the resolution of the post-hoc analyses we focused on the experimental group, in which the majority of fish showed a brief stress response (i.e. the fish exposed to frequent predators).

**Fig. 2:**
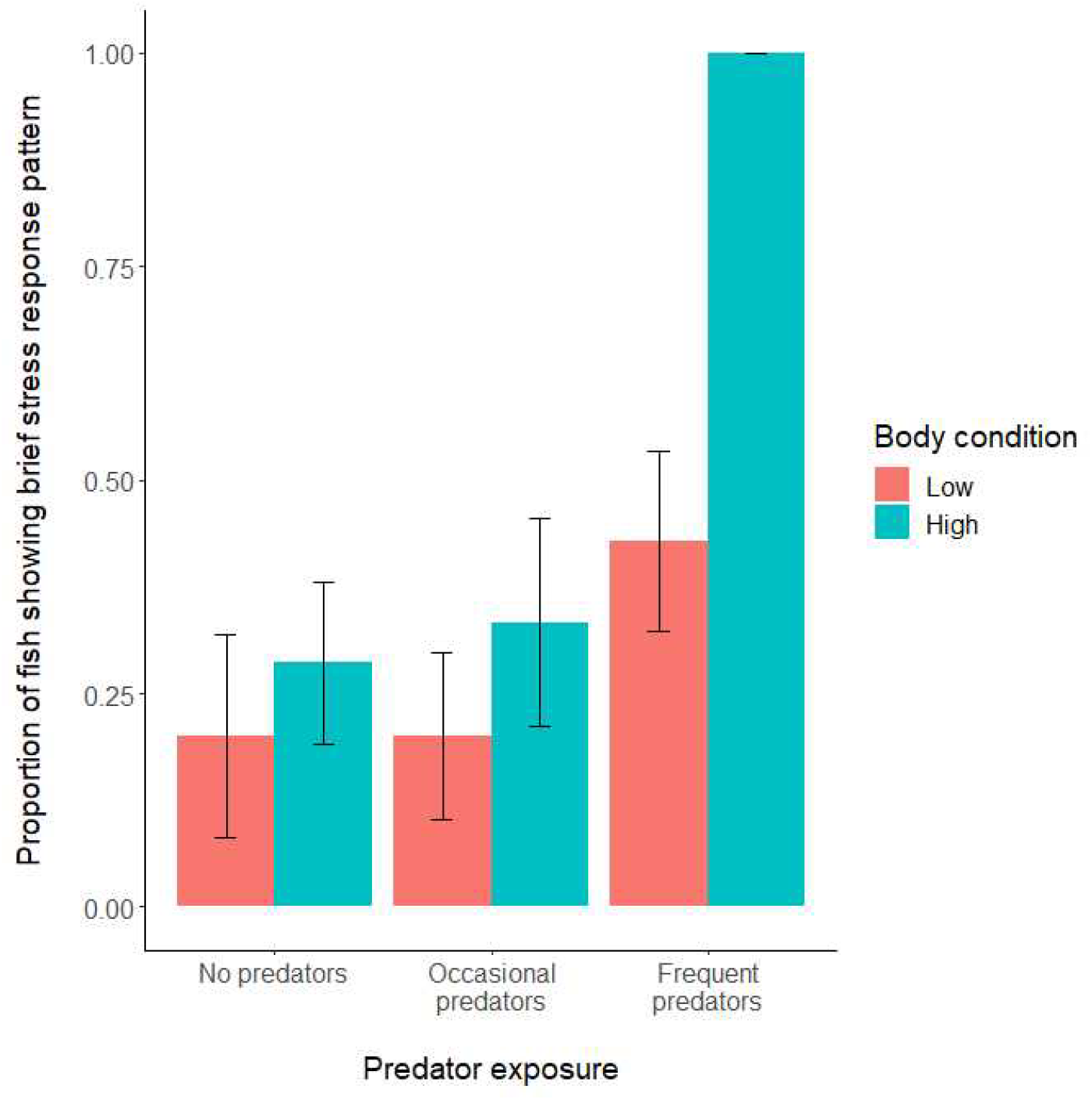
Proportion (mean ± SE) of fish (N=34) showing brief stress response pattern within two hours on the first water sample collection day. Fish were classified as brief or prolonged responders depending on their cortisol concentration profile (see methods for more details). In short brief responders decreased cortisol concentrations within two hours whereas prolonged responders did not. Fish were classified based on their body condition (bc) into: *‘*low body condition*’* (red bars, fish with a bc lower than the average) or *‘*high body condition*’* (blue bars, fish with a bc higher than the average). Groups were exposed to three predator exposures treatments that included the presentation of a control condition without any predators present (*‘*no predators*’*), or the presentations of predators with varying frequencies (*‘*occasional predators*’* or *‘*frequent predators*’*). Fish exposed to the frequent predator treatment were more likely to show a brief stress response pattern compared to fish exposed to occasional predators. For statistical analysis see Table 1 and Supplementary Table S5.

To analyze behavioral flexibility, we used the number of trials until fish reached the learning criterion in the reversal learning phase as the dependent variable and included body condition (*‘*high*’, ‘*low’), predator exposure (‘no predators’, frequent predators’, ‘occasional predators’) as fixed factors in an LMM. To assess whether the treatments influenced the accuracy of fish during the reversal phase, we used the proportion of wrong choices over all trials in this phase (*‘*error rate*’*) (Bannier et al., 2017) as the depend variable and included body condition and predator exposure as fixed effects in a GLMM with a binomial error distribution. To assess whether the treatments influenced the individual’s performance during the acquisition phase, we used the number of trials and the error rate in a LMM and GLMM respectively with body condition and predator exposure as fixed effects.

**Table 1:**
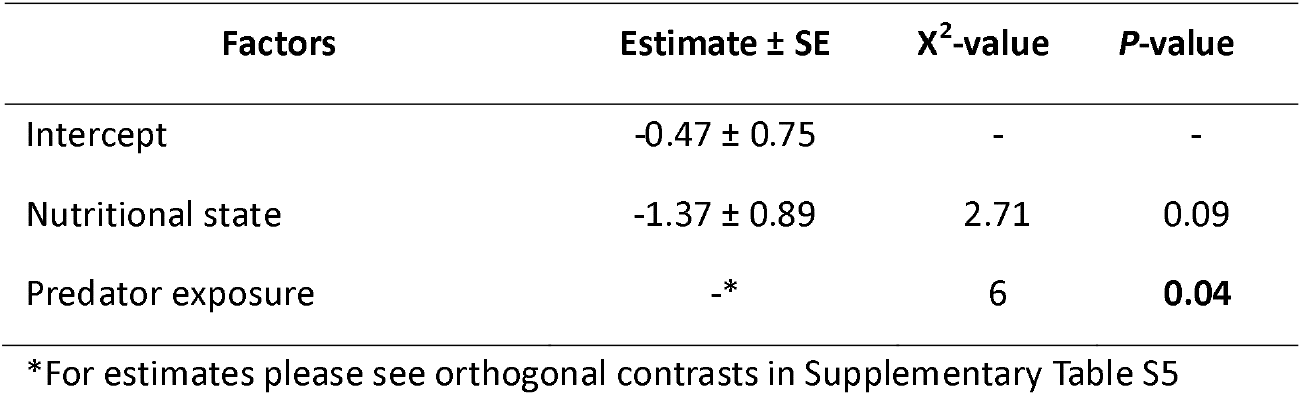
Probability of fish showing acute stress response to a novel stressor. Results are shown from a generalized linear mixed effect model with a binomial error distribution. Predator exposure significantly influenced the probability to show an acute stress response, which was irrespective of the nutritional state. Fish were classified as acute or prolonged responders depending on their cortisol concentration profile for two hours on the first water collection day (see methods for more details). In short acute responders decreased cortisol concentrations within two hours whereas prolonged responders did not. Experimental treatments consisted of the independent manipulations of the subject*’*s nutritional state and predator exposure. Nutritional state was included as a two-level factor (high, low) and the estimate is shown as difference to the reference level (*‘*high*’*). Predator exposure was included as a three-level factor (*‘*no predator*’, ‘*occasional predators*’*, frequent predators*’*) and estimates are shown as orthogonal contrasts in Supplementary Table S5. To obtain p-values a likelihood ration test was used to compare the models with and without the factor of interest. N=34 fish from 20 groups and three blocks. p<0.05 is highlighted in bold.

## Results

### (1) Do nutritional state and the frequency of stressors determine stress recovery patterns?

Contrary to the predictions, the majority of fish exposed to frequent predators had a brief stress response pattern when exposed to a novel stressor, which was not the case for fish exposed to occasional or no predators (Table 1 factor: *‘*Predator exposure [χ2=8.2, p=0.02], for orthogonal contrasts see Supplementary Table S5, Fig. 2). In addition, we found that all fish in a high nutritional state that experienced frequent predators showed a brief stress response pattern (see Fig. 2, N=4 out of 4). To test whether body condition determines stress recovery patterns we performed a post-hoc analysis that used the stress recovery index of fish frequently exposed to predators. Although nutritional status did not predict whether fish were classified as brief or prolonged responders (Table 1 factor: *‘*Body condition*’* [χ2=1.93, p=0.17]), the post-hoc analysis revealed that the duration of the stress response, in terms of the individual stress recovery indices, was clearly influenced by body condition (Supplementary Table S6 factor: *‘*Body condition*’* [F_1,9_=8.78, p=0.02]). Fish with a higher body condition had a more rapid recovery pattern than fish with a lower body condition (see Supplementary Fig. S8). The frequency of predator exposures explained stress response patterns when fish were exposed to a new stressor on day 1 (see Table 1), but not when fish were potentially habituated to the hormone sampling procedures on day 4 (Supplementary Table S7 factor *‘*Predator exposure*’* [χ2=1.48, p=0.48]).

### (2) Do nutritional status and the frequency of stressors determine behavioral flexibility?

In the reversal learning test, fish took on average 27.2 ± 1.5 and 30.6 ± 1.1 (mean ± SEM) trials to reach the learning criterion in the acquisition and reversal phase, respectively. Two fish did not reach the learning criterion in the acquisition phase and were excluded from the analysis. Behavioral flexibility, in terms of the number of trials until fish reached the learning criterion in the reversal phase, varied depending on the frequency of predator exposures and the nutritional status of the fish (Table 2a factor *‘*Body condition x Predator exposure*’* [χ2=5.51, p=0.05]). Contrary to the prediction that frequent stressors and overnutrition would impair behavioral flexibility, orthogonal post-hoc comparisons revealed that fish with high body condition reached the learning criterion faster when they had been exposed to frequent predators compared to fish exposed to occasional predators (Supplementary Table S8c factor *‘*Occasional predators vs. frequent predators (high body condition)*’* [estimate: -0.16, z=-3.4, p<0.01, Fig. 3). Predator exposure did not influence behavioral flexibility of fish with a low body condition (see Supplementary Table S8d,e). Neither predator exposure nor nutritional status influenced the number of trials required to reach the learning criterion in the acquisition phase (Table 2b factors *‘*Body condition*’* [χ^2^=0.05, p=0.83], *‘*Predator exposure*’* [χ^2^=0.14, p=0.93]). Similarly, the experimental manipulations did not affect the error rates of the fish during the acquisition phase (Supplementary Table S9a) and the reversal learning phase (Supplementary Table S9b).

**Table 2:**
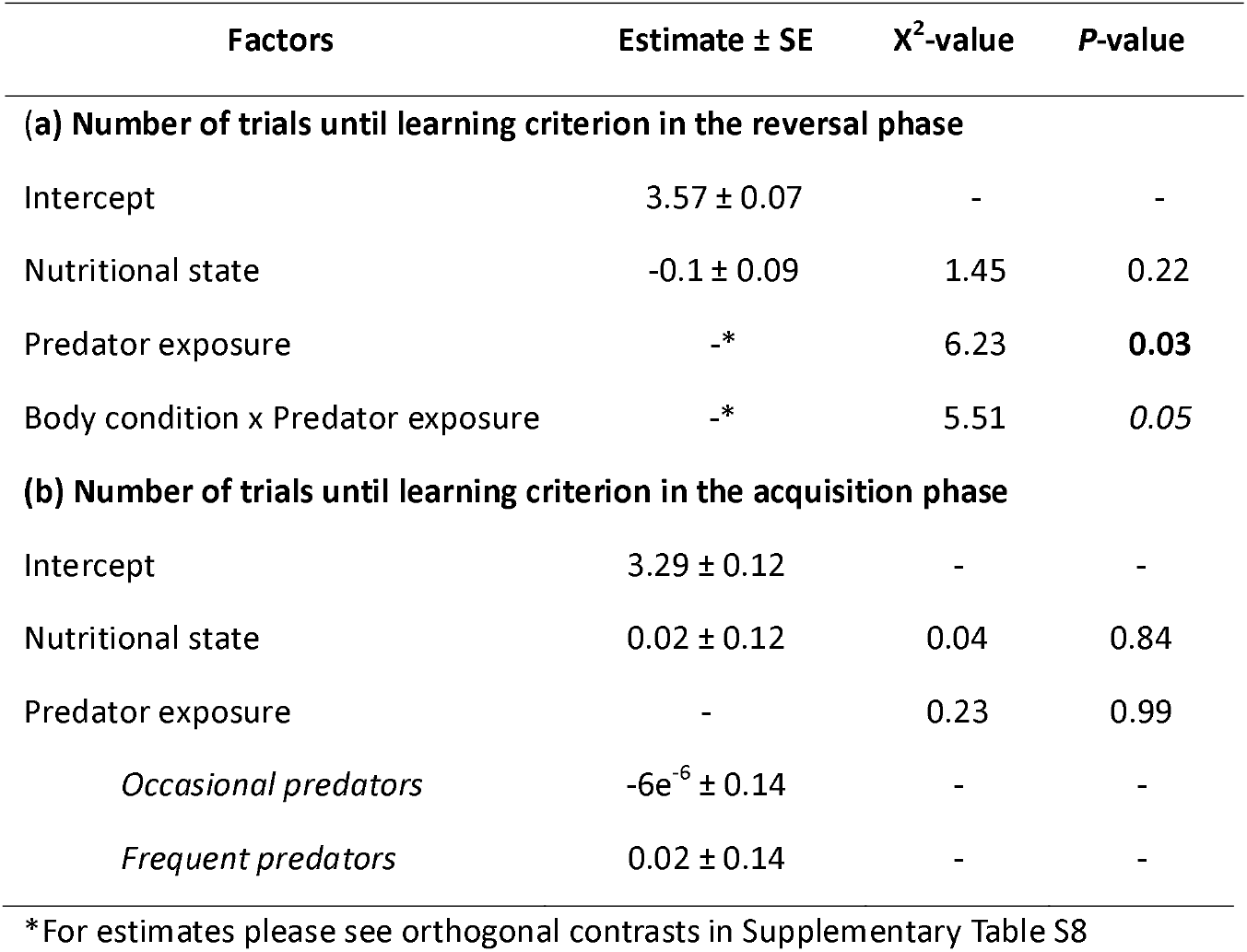
The number of trials until fish reached the learning criterion in the (a) reversal and (b) acquisition phase of the learning task. Results are presented from generalized linear mixed effect models with (a) poisson and (b) negative binomial error distributions. Nutritional state and predator exposure interactively influenced the number of trials until fish reached the learning criterion in the reversal but not in the acquisition phase. During the acquisition phase fish had to discriminate between two colours to access a food reward. During the reversal phase the reward consistency was reversed, and fish were again tested until they associated the new colour with the food reward. Experimental treatments consisted of the independent manipulations of the subject*’*s nutritional state and predator exposure. Nutritional state was included as a two-level factor (high, low) and the estimate is shown as difference to the reference level (*‘*high*’*). Predator exposure was included as a three-level factor (*‘*no predator*’, ‘*occasional predators*’*, frequent predators*’*) and estimates in (a) are shown as orthogonal contrasts in Supplementary Table S8 and in (b) as difference to the reference level *‘*no predators*’*. To obtain p-values a likelihood ration test was used to compare the models with and without the factor of interest. N=34 fish from 20 groups and three blocks. p<0.05 is highlighted in bold and p<0.1 is italicised.

**Fig. 3:**
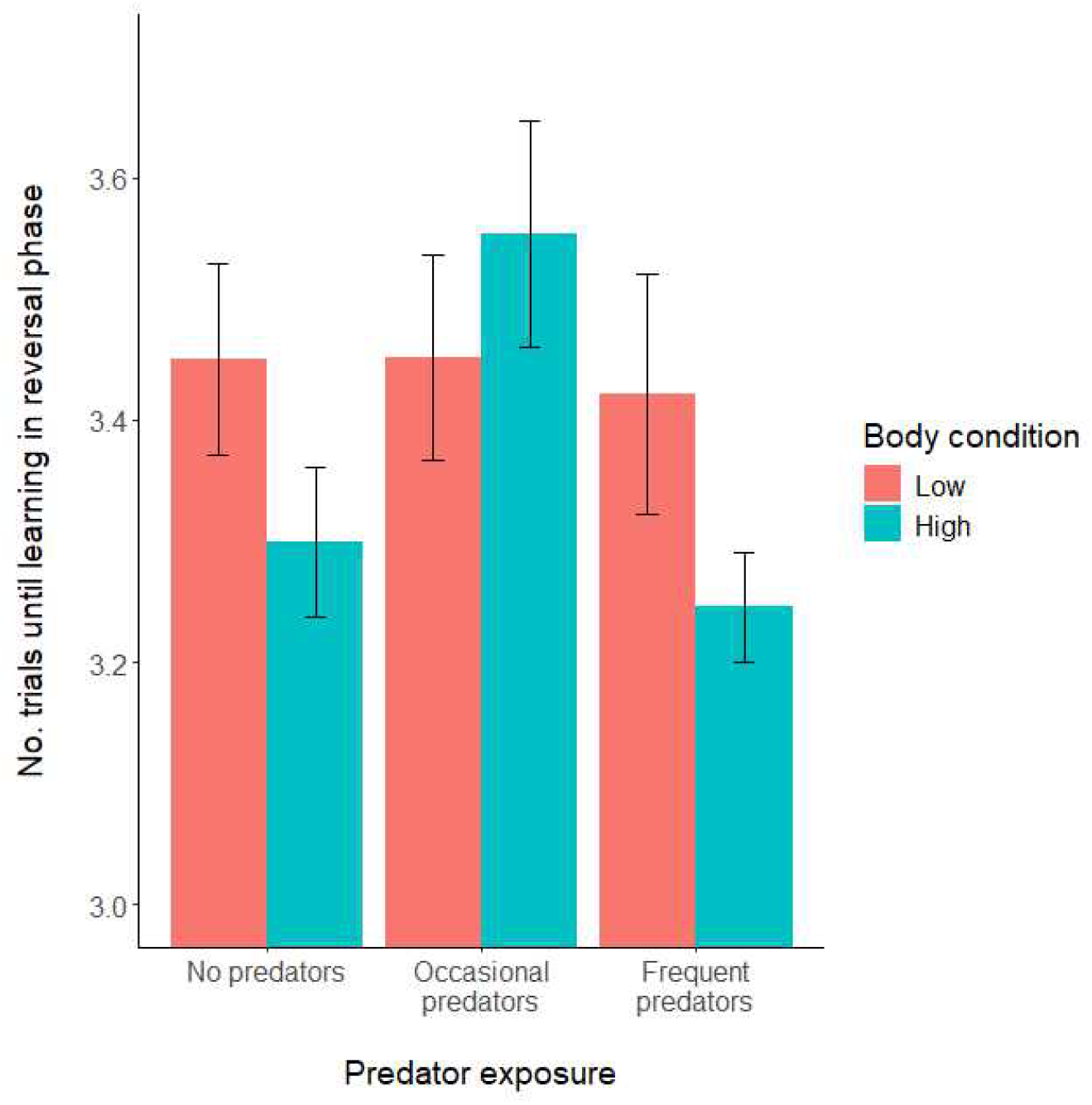
The number of trials (log mean ± SE) until fish reached the learning criterion in the reversal phase of the learning task. Fish were separated into two groups depending on their body condition: low body condition (red bars, lower than the average) or high body condition (blue bars, higher than the average). Groups were exposed to three predator exposure treatments that included the presentation of a control condition without any predators present (*‘*no predators*’*), or the presentations of predators with varying frequencies (*‘*occasional predators*’* or *‘*frequent predators*’*). Fish with a high body condition that were exposed to the frequent predator treatment reached the learning criterion faster than fish with a high body condition that were exposed to the occasional predator treatment. For statistical analysis see Table 2 and Supplementary Table S8

## Discussion

Based on the allostatic load model we predicted that overnutrition and frequent exposure to stressors induce prolonged allostatic states including persistently prolonged stress response patterns and impaired behavioral flexibility. Contrary to this prediction, high body condition fish exposed to frequent predators showed brief stress response patterns when exposed to a new stressor. In addition, high body condition fish exposed to frequent predators also showed greater behavioral flexibility. Therefore, the results do not support our initial hypothesis. An alternative interpretation is that our treatments were insufficient to induce the conditions on which the hypotheses were based. We can think of two reasons for this. (1) Although fish on the increased diet gained more weight and had more energy resources than fish on the reduced diet, they were not obese, with body masses within the adult range (Hirschenhauser et al., 2004), and therefore their condition did not represent overnutrition (McEwen and Wingfield, 2003); (2) The predator presentations clearly induced stress responses in the test fish (Table 1), however, the frequent predator encounters may not have been enough to induce severe chronic stress.

In any case, our results highlight a potential association between brief stress response patterns, characterized by a quick termination of increased cortisol release, and high levels of behavioral flexibility measured as reversal learning speed. This association only became apparent when individuals with high body condition, i. e. with more energy reserves available, were exposed to frequent stressors. However, this association needs to be interpreted with caution due to the low sample size for fish with a high body condition and exposed to frequent predators (N=4). Another caveat to consider when interpreting the results is that we only used one of several parameters to assess stress responses. We chose to focus on the termination of the stress response *a-priori* because the allostatic load model makes very clear predictions about how an excessive food supply hinders the termination of stress responses (McEwen, 2005; McEwen and Wingfield, 2003). Endotherms may be more intuitive models for studying obesity, and under the treatment of the presented study might have resulted in more obese individuals. However, ectotherms, such as fish, can also serve as valuable models to study obesity. For instance, in zebrafish (*Danio rerio*), both overfeeding and high-fat–diet protocols reliably induce increased body mass and adiposity, reproducing key metabolic features of obesity and metabolic syndrome (Ghaddar et al., 2020). In another study, we successfully generated overweight individuals by using a similar feeding protocol (Gabrielidis, 2025). Notably, the overweight fish in the present study exhibited reduced behavioral flexibility and prolonged stress responses, which is consistent with the hypothesis that allostatic load alters neurophysiological and behavioral plasticity.

Glucocorticoid receptors (GR) which are expressed in the brain play an important role in terminating the stress response (de Kloet, 2014; Joels et al., 2008). A previous study that pharmacologically blocked GRs in *N. pulcher* showed that GR blockade reduced behavioral flexibility in a detour task, highlighting a potential link between stress recovery and the ability to cope with changing conditions. Although cortisol was not measured directly, GR-blocked fish also showed more fear-related behaviors, which indicates that blocking GRs led to an accumulation of cortisol due to the reduced ability to terminate the stress response (Fischer et al., 2024). Based on the results of the current study, we can conclude that while resilience (defined as an acute stress response) is correlated with behavioral flexibility, it also incurs associated costs, which well-fed individuals are better able to overcome.

A prolonged phase of allostasis is the result of a chronic exposure to stressors with high energetic costs and consequences for health and fitness (Romero et al., 2009). We did not find support for the prediction that frequent predator exposures and overnutrition will lead to prolonged allostasis. This prediction was derived from the classic version of the allostatic load model which introduced the concept of allostasis as maintaining homeostastis through change (McEwen and Wingfield, 2003). An extension of the classic version, the reactive scope model, predicts that animals with more energy resources will be better able to cope with repeatedly occurring stressors (Romero et al., 2009). This matches with the result that fish with higher body condition had shorter stress response patterns and a high level of behavioral flexibility when exposed to frequent predators. We conclude that the higher energetic resources enabled them to express an adequate response to a new stressor, i.e. a brief stress response pattern.

Interestingly, fish that had both high body condition and were exposed frequently to predators also showed more behavioral flexibility. This suggests that the ability to exhibit a stress response pattern with a brief cortisol peak followed by a timely recovery is associated with higher behavioral flexibility. Stress influences cognition in a complex way that is dependent on the type and duration of the stressor, the individual physiological response type, and the cognitive task being studied (Giovanniello et al., 2023; Sandi and Pinelo-Nava, 2007). Generally, short term stressors are considered to have positive effects on cognitive abilities required to overcome the stressor, whereas long-term or chronic stress is detrimental (Giovanniello et al., 2023; Roozendaal, 2002). Our results provide a valuable contribution to the fields of stress and cognition by highlighting the importance of nutritional status in determining stress responses and behavioral flexibility, i.e. the potential benefits of brief stress response patterns.

There are two mutually exclusive explanations as to why fish adapted to the increased frequency of the stressor with beneficial effects on physiology and flexibility. First, fish exposed to frequent predator presentations might have been habituated to the re-occurring stressor leading to shorter stress responses when exposed to a new stressor and with beneficial effects for behavioral flexibility. We took great care to avoid habituation of fish to the predator presentations by varying the timing of the exposures and adding additional stressors such as chasing with a net or creating a moving shadow. Yet, the fish potentially may have developed habituation simply because they were exposed more frequently to predator presentations. Second, the brief stress responses and high levels of behavioral flexibility could have been an adaptive response to cope with the frequent predator presentations, which would support the idea of preparative effects of glucocorticoid actions (Harris, 2020; Lattin and Romero, 2013; Sapolsky et al., 2000). The *‘*preparative hypothesis*’* suggests that prior glucocorticoid actions modulate the response to a subsequent stressor in a mediating or suppressive way (Sapolsky et al., 2000). Although not yet tested experimentally, this hypothesis has been used to explain seasonal GC variation. According to the preparative hypothesis both, baseline and stress induced GC levels are higher at challenging times of the year (Bauer et al., 2014; Lattin and Romero, 2013). Yet, stressful periods do not necessarily occur at fixed periods of the year and perceived stress may depend on the social status or demands of the individual (Kotrschal et al., 1998a; Kotrschal et al., 1998b). An interesting extension of the preparative hypothesis suggests that GC actions i) prepare to better cope with a subsequent stressor and ii) increase the threshold when certain stimuli are perceived as a stressor (Vera et al., 2017). The observed interaction between body condition and adaptive stress response patterns when exposed to predators is in line with this extension of the hypothesis because the available energy stores affect the ability of organisms to cope with stressors (Landys et al., 2006).

In conclusion, our results show a potential relationship between stress recovery and behavioral flexibility. Groups of fish that terminated stress responses sooner, showed higher levels of behavioral flexibility. This relationship was only seen in groups of fish exposed to frequent predator exposures, which simulated a higher frequency of stressful events. Experiencing more frequent stressful events thus might have prepared fish to better cope with subsequent stressors, particularly in those with greater energy reserves. These results are consistent with the concept of *‘*wear and tear*’*, which focused on the fact that frequent exposure to stressors is costly (Romero et al., 2009) and animals in a better nutritional condition should be better able to cope with repeated stressors due to their higher energy reserves compared to animals in a poor nutritional status.

## Acknowledgments

We would like to thank Claudia Martina and Noémie Liesch for their help during the learning task, Arne Jungwirth for his help with the predator presentations, Martina Krakhofer for help with animal husbandry and Roland Sasse for logistical support. We are particularly grateful to Stefan Graf, whose invaluable support and dedication at all stages of the project was crucial to its success. This study was funded by the Vienna Science and Technology Fund (CS18-042). The text of this manuscript has been improved with the assistance of AI tools, such as ChatGPT and DeepL, to enhance clarity and language quality.

## Authors*’* contributions

S.F.: Conceptualization, data curation, formal analysis, investigation, methodology, project administration, visualization, writing-original draft, writing-review & editing; K.H.: Conceptualization, funding acquisition, writing-review & editing; B.T.: Conceptualization, funding acquisition, methodology, writing-review & editing; L.F.: Conceptualization, funding acquisition, writing-review & editing; V.C.: Investigation, methodology, writing-review & editing; S.T.: Conceptualization, funding acquisition, project administration, supervision, writing-review & editing. (Categories based on CRediT)

## Data accessibility statement

The R-code and the experimental datasets that support the findings of this study are available in Figshare with the identifier https://figshare.com/s/346eb4333a356f0e246b

## Literature

Antunes, D.F., Reyes-Contreras, M., Glauser, G., Taborsky, B., 2021. Early social experience has life-long effects on baseline but not stress-induced cortisol levels in a cooperatively breeding fish. Horm. Behav. 128, 10.1016/j.yhbeh.2020.104910.

ASAB Ethical Committee, 2023. Guidelines for the ethical treatment of nonhuman animals in behavioural research and teaching. Anim. Behav. 195, 10.1016/j.anbehav.2022.09.006.

Balshine, S., Leach, B., Neat, F., Reid, H., Taborsky, M., Werner, N., 2001. Correlates of group size in a cooperatively breeding cichlid fish (Neolamprologus pulcher). Behav. Ecol. Sociobiol. 50, 134–140, 10.1007/s002650100343.

Bannier, F., Tebbich, S., Taborsky, B., 2017. Early experience affects learning performance and neophobia in a cooperatively breeding cichlid. Ethology 123, 712–723, 10.1111/eth.12646.

Bates, D., Maechler, M., Bolker, B.M., Walker, S., 2015. Fitting linear mixed-effects models using {lme4}. J. Stat. Softw. 67, 1–48, 10.18637/jss.v067.i01.

Bauer, C.M., Hayes, L.D., Ebensperger, L.A., Romero, L.M., 2014. Seasonal variation in the degu (Octodon degus) endocrine stress response. Gen. Comp. Endocrinol. 197, 26–32, 10.1016/j.ygcen.2013.11.025.

Benson, S., Arck, P.C., Tan, S., Mann, K., Hahn, S., Janssen, O.E., Schedlowski, M., Elsenbruch, S., 2009. Effects of obesity on neuroendocrine, cardiovascular, and immune cell responses to acute psychosocial stress in premenopausal women. Psychoneuroendocrino. 34, 181–189, 10.1016/j.psyneuen.2008.08.019.

Bryce, C.A., Howland, J.G., 2015. Stress facilitates late reversal learning using a touchscreen-based visual discrimination procedure in male Long Evans rats. Behav. Brain Res. 278, 21–28, 10.1016/j.bbr.2014.09.027.

Buechel, S.D., Boussard, A., Kotrschal, A., van der Bijl, W., Kolm, N., 2018. Brain size affects performance in a reversal-learning test. Proc. R. Soc. B-Biol. Sci. 285, 10.1098/rspb.2017.2031.

Burkart, J.M., Schubiger, M.N., van Schaik, C.P., 2017. The evolution of general intelligence. Behav. Brain Sci. 40, 10.1017/s0140525x16000959.

Canoine, V., Hayden, T.J., Kevin, R., Goymann, W., 2002. The stress response of european stonechats depends on the type of stressor. Behaviour 139, 1303–1311, 10.1163/156853902321104172.

Chbeir, S., Carrión, V., 2023. Resilience by design: How nature, nurture, environment, and microbiome mitigate stress and allostatic load. World J Psychiatry 13, 144–159, 10.5498/wjp.v13.i5.144.

Chida, Y., Steptoe, A., 2010. Greater cardiovascular responses to laboratory mental stress are associated with poor subsequent cardiovascular risk status: a meta-analysis of prospective evidence. Hypertension 55, 1026–1032, 10.1161/hypertensionaha.109.146621.

Creel, S., Dantzer, B., Goymann, W., Rubenstein, D.R., 2013. The ecology of stress: effects of the social environment. Funct. Ecol. 27, 66–80, 10.1111/j.1365-2435.2012.02029.x.

D’Amico, D., Amestoy, M.E., Fiocco, A.J., 2020. The association between allostatic load and cognitive function: a systematic and meta-analytic review. Psychoneuroendocrino. 121, 104849, 10.1016/j.psyneuen.2020.104849.

de Kloet, E.R., 2014. From receptor balance to rational glucocorticoid therapy. Endocrinology 155, 2754–2769, 10.1210/en.2014-1048.

de Kloet, E.R., Joels, M., Holsboer, F., 2005. Stress and the brain: from adaptation to disease. Nat. Rev. Neurosci. 6, 463–475, 10.1038/nrn1683.

Diamond, A., 2013. Executive functions, in: Fiske, S.T. (Ed.), Annual Review of Psychology, Vol 64. Annual Reviews, Palo Alto, pp. 135–168.

Dickens, M.J., Romero, L.M., 2013. A consensus endocrine profile for chronically stressed wild animals does not exist. Gen Comp Endocrinol 191, 177–189, 10.1016/j.ygcen.2013.06.014.

Engqvist, L., 2005. The mistreatment of covariate interaction terms in linear model analyses of behavioural and evolutionary ecology studies. Anim. Behav. 70, 967–971, 10.1016/j.anbehav.2005.01.016.

Fischer, S., Balshine, S., Hadolt, M.C., Schaedelin, F.C., 2021. Siblings matter: Family heterogeneity improves associative learning later in life. Ethology 127, 897–907, 10.1111/eth.13196.

Fischer, S., Ferlinc, Z., Hirschenhauser, K., Taborsky, B., Fusani, L., Tebbich, S., 2024. Does the stress axis mediate behavioural flexibility in a social cichlid, Neolamprologus pulcher? Physiol. Behav. 287, 114694, 10.1016/j.physbeh.2024.114694.

Gabrielidis, X.-N., 2025. The influence of the nutritional state on stress responses and behavioural flexibility. Masterthesis, University of Vienna, 10.25365/thesis.79412.

George, S.A., Rodriguez-Santiago, M., Riley, J., Abelson, J.L., Floresco, S.B., Liberzon, I., 2015. Alterations in cognitive flexibility in a rat model of post-traumatic stress disorder. Behav. Brain Res. 286, 256–264, 10.1016/j.bbr.2015.02.051.

Ghaddar, B., Veeren, B., Rondeau, P., Bringart, M., Lefebvre d’Hellencourt, C., Meilhac, O., Bascands, J.-L., Diotel, N., 2020. Impaired brain homeostasis and neurogenesis in diet-induced overweight zebrafish: a preventive role from A. borbonica extract. Sci. Rep. 10, 14496, 10.1038/s41598-020-71402-2.

Giovanniello, J., Bravo-Rivera, C., Rosenkranz, A., Matthew Lattal, K., 2023. Stress, associative learning, and decision-making. Neurobiol Learn Mem 204, 107812, 10.1016/j.nlm.2023.107812.

Graybeal, C., Feyder, M., Schulman, E., Saksida, L.M., Bussey, T.J., Brigman, J.L., Holmes, A., 2011. Paradoxical reversal learning enhancement by stress or prefrontal cortical damage: rescue with BDNF. Nat. Neurosci. 14, 1507–1509, 10.1038/nn.2954.

Groenewoud, F., Frommen, J.G., Josi, D., Tanaka, H., Jungwirth, A., Taborsky, M., 2016. Predation risk drives social complexity in cooperative breeders. Proc. Natl. Acad. Sci. U.S.A., 4104–4109, 10.1073/pnas.1524178113.

Harris, B.N., 2020. Stress hypothesis overload: 131 hypotheses exploring the role of stress in tradeoffs, transitions, and health. Gen. Comp. Endocrinol. 288, 113355, 10.1016/j.ygcen.2019.113355.

Heg, D., Brouwer, L., Bachar, Z., Taborsky, M., 2005. Large group size yields group stability in the cooperatively breeding cichlid Neolamprologus pulcher. Behaviour 142, 1615–1641, 10.1163/156853905774831891.

Hirschenhauser, K., Taborsky, M., Oliveira, T., Canario, A.V.M., Oliveira, R.F., 2004. A test of the ‘challenge hypothesis’ in cichlid fish: simulated partner and territory intruder experiments. Anim. Behav. 68, 741–750, 10.1016/j.anbehav.2003.12.015.

Joels, M., Karst, H., DeRijk, R., de Kloet, E.R., 2008. The coming out of the brain mineralocorticoid receptor. Trends Neurosci. 31, 1–7, 10.1016/j.tins.2007.10.005.

Jones, A., McMillan, M.R., Jones, R.W., Kowalik, G.T., Steeden, J.A., Deanfield, J.E., Pruessner, J.C., Taylor, A.M., Muthurangu, V., 2012. Adiposity is associated with blunted cardiovascular, neuroendocrine and cognitive responses to acute mental stress. Plos ONE 7, e39143, 10.1371/journal.pone.0039143.

Jordan, A., Taborsky, B., Taborsky, M., 2021. Cichlids as a Model System for Studying Social Behaviour and Evolution, in: Abate, M.E., Noakes, D.L.G. (Eds.), The Behavior, Ecology and Evolution of Cichlid Fishes. Springer Netherlands, Dordrecht, pp. 587–635.

Jungwirth, A., Balzarini, V., Zottl, M., Salzmann, A., Taborsky, M., Frommen, J.G., 2019. Long-term individual marking of small freshwater fish: the utility of Visual Implant Elastomer tags. Behav. Ecol. Sociobiol. 73, 10.1007/s00265-019-2659-y.

Kitaysky, A.S., Piatt, J.F., Wingfield, J.C., Romano, M., 1999. The adrenocortical stress-response of Black-legged Kittiwake chicks in relation to dietary restrictions. Journal of Comparative Physiology B 169, 303–310, 10.1007/s003600050225.

Kotrschal, K., Hirschenhauser, K., Möstl, E., 1998a. The relationship between social stress and dominance is seasonal in greylag geese. Anim. Behav. 55, 171–176, 10.1006/anbe.1997.0597.

Kotrschal, K., Van Staaden, M.J., Huber, R., 1998b. Fish brains: evolution and environmental relationships. Reviews in Fish Biology and Fisheries 8, 373–408, 10.1023/a:1008839605380.

Kühnel, A., Hagenberg, J., Knauer-Arloth, J., Ködel, M., Czisch, M., Sämann, P.G., Binder, E.B., Kroemer, N.B., 2023. Stress-induced brain responses are associated with BMI in women. Commun Biol 6, 1031, 10.1038/s42003-023-05396-8.

Landys, M.M., Ramenofsky, M., Wingfield, J.C., 2006. Actions of glucocorticoids at a seasonal baseline as compared to stress-related levels in the regulation of periodic life processes. Gen Comp Endocrinol 148, 132–149, 10.1016/j.ygcen.2006.02.013.

Lattin, C.R., Romero, L.M., 2013. Seasonal variation in corticosterone receptor binding in brain, hippocampus, and gonads in house sparrows (Passer domesticus). The Auk 130, 591–598, 10.1525/auk.2013.13043.

Lynn, S.E., Hunt, K.E., Wingfield, J.C., 2003. Ecological factors affecting the adrenocortical response to stress in chestnut-collared and McCown’s longspurs (Calcarius ornatus, Calcarius mccownii). Physiol Biochem Zool 76, 566–576, 10.1086/375435.

MacLeod, K.J., English, S., Ruuskanen, S.K., Taborsky, B., 2023. Stress in the social context: a behavioural and eco-evolutionary perspective. J. Exp. Biol. 226, 10.1242/jeb.245829.

McEwen, B.S., 2005. Stressed or stressed out: what is the difference? J. Psychiatr. Neurosci. 30, 315–318.

McEwen, B.S., Wingfield, J.C., 2003. The concept of allostasis in biology and biomedicine. Horm. Behav. 43, 2–15, 10.1016/s0018-506x(02)00024-7.

McEwen, B.S., Wingfield, J.C., 2010. What is in a name? Integrating homeostasis, allostasis and stress. Horm Behav 57, 105–111, 10.1016/j.yhbeh.2009.09.011.

Mujica-Parodi, L.R., Renelique, R., Taylor, M.K., 2009. Higher body fat percentage is associated with increased cortisol reactivity and impaired cognitive resilience in response to acute emotional stress. Int J Obes (Lond) 33, 157–165, 10.1038/ijo.2008.218.

Nyman, C., Fischer, S., Aubin-Horth, N., Taborsky, B., 2017. Effect of the early social environment on behavioural and genomic responses to a social challenge in a cooperatively breeding vertebrate. Mol. Ecol. 26, 3186–3203, 10.1111/mec.14113.

Nyman, C., Fischer, S., Aubin-Horth, N., Taborsky, B., 2018. Evolutionary conserved neural signature of early life stress affects animal social competence. Proc. R. Soc. B-Biol. Sci. 285, 10.1098/rspb.2017.2344.

Ochi, H., Yanagisawa, Y., 1998. Commensalism between cichlid fishes through differential tolerance of guarding parents toward intruders. J. Fish Biol. 52, 985–996, 10.1111/j.1095-8649.1998.tb00598.x.

Petry, N.M., Barry, D., Pietrzak, R.H., Wagner, J.A., 2008. Overweight and obesity are associated with psychiatric disorders: results from the National Epidemiologic Survey on Alcohol and Related Conditions. Psychosom Med 70, 288–297, 10.1097/PSY.0b013e3181651651.

R Core Team, 2024. R: a language and environment for statistical computing. R Foundation for Statistical Computing, Vienna, Austria.

Reyes-Contreras, M., Glauser, G., Rennison, D.J., Taborsky, B., 2019. Early-life manipulation of cortisol and its receptor alters stress axis programming and social competence. Philos. Trans. R. Soc. B-Biol. Sci. 374, 10.1098/rstb.2018.0119.

Reyes-Contreras, M., Taborsky, B., 2022. Stress axis programming generates long-term effects on cognitive abilities in a cooperative breeder. Proc. R. Soc. B-Biol. Sci. 289, 10.1098/rspb.2022.0117.

Romero, L.M., Dickens, M.J., Cyr, N.E., 2009. The reactive scope model - a new model integrating homeostasis, allostasis, and stress. Horm. Behav. 55, 375–389, 10.1016/j.yhbeh.2008.12.009.

Roozendaal, B., 2002. Stress and memory: opposing effects of glucocorticoids on memory consolidation and memory retrieval. Neurobiol Learn Mem 78, 578–595, 10.1006/nlme.2002.4080.

Sandi, C., Pinelo-Nava, M.T., 2007. Stress and memory: behavioral effects and neurobiological mechanisms. Neural Plasticity 2007, 10.1155/2007/78970.

Sapolsky, R.M., Romero, L.M., Munck, A.U., 2000. How do glucocorticoids influence stress responses? Integrating permissive, suppressive, stimulatory, and preparative actions. Endocr. Rev. 21, 55–89, 10.1210/er.21.1.55.

Schultner, J., Kitaysky, A.S., Welcker, J., Hatch, S., 2013. Fat or lean: adjustment of endogenous energy stores to predictable and unpredictable changes in allostatic load. Funct. Ecol. 27, 45–55, 10.1111/j.1365-2435.2012.02058.x.

Schwabe, L., Joels, M., Roozendaal, B., Wolf, O.T., Oitzl, M.S., 2012. Stress effects on memory: an update and integration. Neurosci. Biobehav. Rev. 36, 1740–1749, 10.1016/j.neubiorev.2011.07.002.

Scott, A.P., Hirschenhauser, K., Bender, N., Oliveira, R., Earley, R.L., Sebire, M., Ellis, T., Pavlidis, M., Hubbard, P.C., Huertas, M., Canario, A., 2008. Non-invasive measurement of steroids in fish-holding water: important considerations when applying the procedure to behaviour studies. Behaviour 145, 1307–1328, 10.1163/156853908785765854.

Singmann, H., Bolker, B., Westfall, J., Aust, F., Ben-Shachar, M.S., 2021. afex: analysis of factorial experiments, R package version 1.0-1 ed.

Sol, D., Timmermans, S., Lefebvre, L., 2002. Behavioural flexibility and invasion success in birds. Anim. Behav. 63, 495–502, 10.1006/anbe.2001.1953.

Spencer, S.J., Tilbrook, A., 2009. Neonatal overfeeding alters adult anxiety and stress responsiveness. Psychoneuroendocrino. 34, 1133–1143, 10.1016/j.psyneuen.2009.02.013.

Taborsky, B., Tschirren, L., Meunier, C., Aubin-Horth, N., 2013. Stable reprogramming of brain transcription profiles by the early social environment in a cooperatively breeding fish. Proceedings of the Royal Society of London Series B-Biological Sciences 280, 10.1098/rspb.2012.2605.

Taborsky, M., 1984. Broodcare helpers in the cichlid fish Lamprologus brichardi: their costs and benefits. Anim. Behav. 32, 1236–1252, 10.1016/s0003-3472(84)80241-9.

Taborsky, M., 1985. Breeder-helper conflict in a cichlid fish with broodcare helpers: an experimental analysis. Behaviour 95, 45–75, 10.1163/156853985x00046.

Tebbich, S., Sterelny, K., Teschke, I., 2010. The tale of the finch: adaptive radiation and behavioural flexibility. Philos. Trans. R. Soc. B-Biol. Sci. 365, 1099–1109, 10.1098/rstb.2009.0291.

Thai, C.A., Zhang, Y., Howland, J.G., 2013. Effects of acute restraint stress on set-shifting and reversal learning in male rats. Cogn. Affect. Behav. Neurosci. 13, 164–173, 10.3758/s13415-012-0124-8.

Vera, F., Zenuto, R., Antenucci, C.D., 2017. Expanding the actions of cortisol and corticosterone in wild vertebrates: a necessary step to overcome the emerging challenges. Gen Comp Endocrinol 246, 337–353, 10.1016/j.ygcen.2017.01.010.

Watve, M., Prati, S., Taborsky, B., 2019. Simulating more realistic predation threat using attack playbacks. PeerJ 7, 16, 10.7717/peerj.8149.

Wingfield, J.C., 2013. The comparative biology of environmental stress: behavioural endocrinology and variation in ability to cope with novel, changing environments. Anim. Behav. 85, 1127–1133, 10.1016/j.anbehav.2013.02.018.

Wong, S.C., Dykstra, M., Campbell, J.M., Earley, R.L., 2008. Measuring water-borne cortisol in convict cichlids (Amatitlania nigrofasciata): is the procedure a stressor? Behaviour 145, 1283–1305, 10.1163/156853908785765863.

Wright, J., Buch, K., Beattie, U.K., Gormally, B.M.G., Romero, L.M., Fefferman, N., 2023. A mathematical representation of the reactive scope model. Journal of Mathematical Biology 87, 51, 10.1007/s00285-023-01983-9.

