## Supplemental Information for "Fat fish stay cool: Stress recovery and behavioral flexibility vary with nutritional state and predator exposure in the cichlid *Neolamprologus pulcher*"

Electronic supplementary material for:

The following Supplementary Material includes additional methodological details Supplementary methods S1, S3, S4, S5), a description of a pilot study (Supplementary Methods S2, Supplementary Tables S1, S2 and Supplementary Figs. S4, S5), details on each of the statistical models presented in the Results section (Supplementary Table S3), additional tables (Supplementary Tables S4-S9) and figures (Supplementary Figs. S6-S8) to support the conclusions presented in the main body of the manuscript, and three figures (Supplementary Figs. S1-S3) providing further details on the experimental design and hormone sampling procedure.

### Supplementary Methods S1

Prior to the start of the experiment, we measured the standard length of the test fish (from the tip of the snout to the end of the caudal peduncle [SL], to the nearest 0.1 cm) and body mass (BM, to the nearest 0.1 g). To obtain individual growth rates (see Supplementary Figs. S6, S7) and to avoid testing the same fish repeatedly, we used visible implant elastomer (VIE, Northwest Marine Technology, Inc.) tags to individually mark the fish [1]. In block 1, all test fish were divided into six groups of seven, separated by sex (three female and three male groups) and placed in 160L tanks. In blocks 2 and 3, test fish showed more signs of aggression prior to the start of the experiment, and we decided, for ethical reasons, to separate the fish into 12 groups of three (six male and six female groups) and 12 groups of four (six male and six female groups) again separated by sex (see Supplementary Fig. S1).

### Supplementary Methods S2

We conducted a pilot study consisting of two phases to investigate (1) whether the increased and reduced feeding regime create a detectable change in body condition and (2) how long each group would take to recover from their respective previous diet and show an overlapping body condition (BC). To do so, we randomly selected nine adult *N pulcher* from the stock population that were not part of the main experiment. In the first phase, we initially measured their body mass and standard length and randomly assigned five individuals to the increased (1.5 sticks per fish of JBL Stick M for cichlids©) and four to the reduced (0.5 sticks per fish of JBL Stick M for cichlids©) diet. In both feeding regimes the fish were fed in the same way as described in the main experiment six times per week for nine weeks. During this time, we repeatedly measured their body mass and body size on a weekly basis. To analyse whether the feeding regimes influenced the body conditions (BC) of pilot fish we calculated BC using the formula: $BC=(({\frac{BM}{SL})}^{3})\times100$ for each measurement time point. Then, to analyse whether the individual BCs changed throughout the feeding regime, we calculated the difference of each measurement taken during the feeding regime to the measurement taken before the pilot experiment started. We analysed this difference using a linear mixed effect model with the type of diet (increased, reduced) as a fixed factor, the time of measurement as a covariate and the individual identity as a random effect. The results show that the two feeding regimes had the desired effect. Fish on the increased diet showed a consistently higher BC compared to fish on the reduced diet (see factor “Diet” [F_1,7_=13.81, p=0.01] in Supplementary Table S1, Fig. S4).

In the second phase of the pilot experiment we switched each group of fish to an intermediate diet (1 stick per fish of JBL Stick M for cichlids^©^) for another five weeks. During this time we continued the weekly measurements of individual fish to detect the time point when BCs of the two groups start to overlap. The change of BC was again calculated as the difference of the measurements taken during the second phase of the pilot experiment and the measurement taken before the feeding started in the first phase. We analysed the change in BC using a linear mixed effect model with the direction of the diet change (increased to intermediate, reduced to intermediate) as a fixed effect, the measurement time as a covariate, and the individual identity as a random effect. We found that even after five weeks on the same feeding regime the BCs of the two groups did not entirely overlap. Fish that previously were on an increased diet had higher BCs during the entire course of the second phase of the pilot experiment than fish that previously were on a reduced diet (see factor “Diet change” [F_1,7_=8.11, p=0.02] in Supplementary Table S2, Fig. S5). Thus, the different body conditions resulting from the diet manipulations was enduring for at least five weeks.

### Supplementary Methods S3

To ensure that all groups received predator presentations at different times of the day we allocated presentations on a weekly basis in the way that no more than two presentations for the same group occurred within the same 1h period (for example from 10am -11am). Within this constraint we then randomly selected the time slots of the predator presentations for each group. In addition, to induce more natural escape responses, that are less frequent when using videos instead of live predators (S.F. pers. obs.), we also randomly performed one of three standardized stress manipulations during the predator presentations which included (1) no manipulation, (2) moving of an dark-grey, opaque plate (48 x 35 cm) along the long side of the tank approximately 10 cm above the tank lid for 2s or (3) moving of a hand net (15 x 13 cm) along the long side of the tank inside the tank water for two seconds. Control recordings without any predators were presented three time per week following the same semi-random schedule as described above, without any stress manipulations.

### Supplementary Methods S4

Cortisol in fish water derives from passive diffusion across the gill epithelia [2], as well as urine excretion [3]. Waterborne cortisol is a reliable proxy for circulating plasma cortisol, because (i) cortisol levels derived from holding water correlate positively with plasma cortisol levels [2] and (ii) the lag between cortisol peaks in plasma and water is only a few minutes [4]. A water flow-through system was established to collect several, consecutive holding water samples from the same individual without disturbing the fish. The system consisted of a 45L tank serving as clean tap water reservoir, two 1-L fish-holding containers and two 1-L Erlenmeyer flasks to collect the hormone-water samples. Water with excreted cortisol from the fish-holding containers accumulated in the collecting flasks at a constant rate, while the fish-holding container was constantly refilled by clean tap water from the reservoir at the same rate. The collecting flasks were exchanged every 30 minutes, and the 500 ml sample water was immediately transferred to a -20°C freezer until further processing. This resulted into four water samples throughout the 2h sampling period. Water flow was regulated in a way that there was one complete exchange of the water volume of the fish-holding container while collecting one sample (30 min). Two samples were collected in parallel, using a multichannel ISMATEC peristaltic pump (ISM404B, ISMATEC, Switzerland). Teflon (PFTE) tubing (Deutsch & Neumann™) of 4 mm inner diameter was used to pump water between the different compartments. The reservoir tank was never completely depleted and new tap water was always filled in at least one day before the next hormone sampling to ensure that the water in the reservoir tank had always the same temperature and oxygen saturation levels. For this we placed a heater and a biological filter in the reservoir water tank, which were not in contact with any other aquarium water before.

### Supplementary Methods S5

Conditioned cartridges were defrosted, then placed on a vacuum manifold (Biotage VacMaster 20) and eluted with 5ml ethyl acetate (HPLC -grade 99.9%). Ethyl acetate proved to be a very satisfactory solvent for the elution of cortisol (mainly free CORT) from solid phase extraction cartridges [2]. Collected cortisol samples were dried down with N_2_ at 45 °C, resuspend with 750ul assay buffer, vortexed and placed overnight in a fridge. Free cortisol concentrations were quantified using a commercial enzymatic immunoassay for cortisol (ENZO Life Sciences Cat # ADI 900-071). Reconstituted samples were brought to RT on a shaker for 30 min and then further processed following the manufacture’s instruction. All samples were analysed in duplicates. The sensitivity of the assay was 24.7 pg/ml, intra-assay CV% was ≤ 5% and inter assay CV% was below 13% using a control sample. Sample concentrations were corrected for dilution. Of the 36 sampled test fish, we were not able to reliably measure cortisol from 2 fish in any of the collected water samples, reducing the overall sample size to 34 fish.

### Supplementary Table S1

| **Factors** | **Estimate ± SE** | **Num. D.F.** | **Den. D.F** | **F-value** | ***P*-value** |
| --- | --- | --- | --- | --- | --- |
| Intercept | 0.18 ± 0.09 |  |  |  |  |
| Diet | -0.38 ± 0.1 | 1 | 7 | 13.81 | **0.01** |
| Measurement time point | 0.04 ± 0.01 | 1 | 35 | 9.24 | **<0.01** |

**Table S1**: The change in body condition of fish on two different diets (reduced, increased) during the first phase of the pilot experiment. The change in body condition (dependent variable) was calculated for each individual as the difference between the body condition measured on a specific time point during the feeding regime (one week, two weeks, three weeks, four weeks, five weeks, six weeks, eight weeks, and nine weeks on their respective diets, see Fig. S4) and the body condition before the feeding started. Over the course of the first phase of the pilot experiment, fish on the increased diet developed a higher body condition than fish on the reduced diet (see also Fig. S4). Results are shown from a linear mixed model. The estimate for factor ‘Diet’ is shown as differences to the reference level ‘increased’. To obtain p-values an F-test was used to compare a model with the factor of interest to a model without it. N=9 individuals in 45 observations.

### Supplementary Table S2

| **Factors** | **Estimate ± SE** | **Num. D.F.** | **Den. D.F** | **F-value** | ***P*-value** |
| --- | --- | --- | --- | --- | --- |
| Intercept | 0.41 ± 0.08 | - | - | - | - |
| Diet change | -0.3 ± 0.11 | 1 | 7 | 8.11 | **0.02** |
| Measurement time point | -0.02 ± 0.02 | 1 | 26 | 0.76 | 0.39 |

**Table S2:** The change in body condition of fish after the switch to an intermediate diet during the second phase of the pilot experiment. The change in body condition (dependent variable) was calculated for each individual as the difference between the body condition measured on a specific time point after the switch in feeding regime (one week, two weeks, three weeks, four weeks, five weeks on the intermediate diet, see Fig. S5) and the body condition before the previous feeding regime. Over the course of the second phase of the pilot experiment fish that were previously on an increased diet had still significantly higher BCs than fish that were previously on a reduced diet (see also Fig. S5). Results are shown from a linear mixed model. The estimate for factor ‘Diet change’ is shown as differences to the reference level ‘increased to intermediate’. To obtain p-values an F-test was used to compare a model with the factor of interest to a model without it. N=9 individuals in 36 observations.

### Supplementary Table S3.

| **Response variable** | **Explanatory variables** | **Covariates** | **Random factor** | **Transformation or Link** | **Interactions** | **Final model** |
| --- | --- | --- | --- | --- | --- | --- |
| Whether fish showed an acute (1) or a prolonged (0) stress response on the first water sample collection day | Nutritional state  Predator exposure | Sex | Block ID  Tank ID | Logit link  (GLMM) | Nutritional state x Predator exposure | Table 1 |
| Intensity of stress reactivity | Body condition | Sex | Block ID  Tank ID | Log  (LMM) | None | Table 2 |
| Nr. of trials until fish reached learning criterion in the reversal phase | Nutritional state  Predator exposure | Sex | Block ID  Tank ID | Log  (GLMM) | Nutritional state x Predator exposure | Table 3a |
| Nr. of trials until fish reached learning criterion in the acquisition phase | Nutritional state  Predator exposure | Sex | Block ID  Tank ID | Log  (GLMM) | Nutritional state x Predator exposure | Table 3a |
| Change in body condition | Diet | Measurement time point | Individual ID | None  (LMM) | None | Table S1 |
| Change in body condition | Diet change | Measurement time point | Individual ID | None  (LMM) | None | Table S2 |
| Daily weight gain | Feeding regime  Predator exposure | Sex | Block ID  Tank ID | None  (LMM) | None | Table S4a |
| Daily growth | Feeding regime  Predator exposure | Sex | Block ID  Tank ID | None  (LMM) | None | Table S4b |
| Whether fish showed an acute (1) or a prolonged (0) stress response on the fourth water sample collection day | Nutritional state  Predator exposure | Sex | Block ID  Tank ID | Logit link  (GLMM) | Nutritional state x Predator exposure | Table S6 |
| Error rates during the acquisition phase | Nutritional state  Predator exposure | Sex | Block ID  Tank ID | Logit link  (GLMM) | Nutritional state x Predator exposure | Table S8a |
| Error rates during the reversal phase | Nutritional state  Predator exposure | Sex | Block ID  Tank ID | Logit link  (GLMM) | Nutritional state x Predator exposure | Table S8b |

**Table S3:** Information on the linear mixed models (LMM) and a generalized linear mixed model (GLMM) analysed during this study, including the respective response variables, explanatory variables, covariates and random factors, eventual data transformations performed to obtain normally distributed residuals, and any interaction terms included in the initial, full models. If data included zeros and log transformations had to be applied, we added a constant value to all data points (i.e. 1, or in cases with data points below 1, the value closest to 0). To obtain final models, non-significant covariates and interactions were stepwise removed. Explanations of response variables: ‘Whether fish showed an acute (1) or a prolonged (0) stress response on the first water sample collection day’: Fish were classified as acute or prolonged responders depending on their stress response patterns on the first water sample collection day. Acute stress responders showed a decline whereas prolonged responders did not show a decline in cortisol concentrations within 2h (for more details see methods); ‘Intensity of stress reactivity’: The intensity was calculated as the difference between the individual peak concentration and the final value after 2h on the first water sample collection day for fish exposed to frequent predators (for more details see methods). ‘Nr. of trials until fish reached learning criterion in the reversal phase’ Number of trials until fish reached the learning criterion in the reversal phase of the learning task; ‘Nr. of trials until fish reached learning criterion in the acquisition phase’ Number of trials until fish reached the learning criterion in the acquisition phase of the learning task; ‘Change in body condition’: The change in body condition of separate fish used in a pilot study as the difference between the body condition measured on a specific time point during the pilot study and the body condition before the pilot study started; ‘Whether fish showed an acute (1) or a prolonged (0) stress response on the fourth water sample collection day’: Fish were classified as acute or prolonged responders depending on their stress response patterns on the fourth water sample collection day; ‘Error rates during the acquisition phase’: Overall error rates, calculated as number of error divided by the overall number of trials of fish during the acquisition phase of the learning task; ‘Error rates during the reversal phase’: Overall error rates, calculated as number of error divided by the overall number of trials of fish during the reversal phase of the learning task; ‘Daily weight gain’: The individual percentage of daily weight gain of the fish during the study; ‘Daily growth’: The individua percentage of daily growth rates of the fish during the study. Explanatory variable names: ‘Nutritional state’: Fish were either classified as ‘high nutritional state’ or ‘low nutritional state’ depending on whether their body condition was above or below the average; ‘Predator exposure’: Fish were exposed to three frequencies of predator exposures (none, occasional and frequent); ‘Body condition’: The body condition of fish exposed to frequent predators. Covariate names: ‘Sex’: The sex of test fish; ‘Measurement time point’: The time point (in weeks) when fish were measured during a pilot study. Random factor names: ‘Block ID’: identification number of block 1-3; ‘Tank ID’: identification number of the housing tank to control for similar; ‘Individual ID’: individual identification number.

### Supplementary Table S4

| **Factors** | **Estimate ± SE** | **Num. D.F.** | **Den. D.F** | **F-value** | ***P*-value** |
| --- | --- | --- | --- | --- | --- |
| **(a) Daily weight gain of fish during the experiment** | | | | | |
| Intercept | 0.08 ± 0.02 | - | - | - | - |
| Feeding regime | -0.18 ± 0.02 | 1 | 23.9 | 80.7 | **<0.01** |
| Predator exposure | - | 2 | 18.24 | 1.08 | 0.36 |
| Occasional predators | 0.03 ± 0.02 | - | - | - | - |
| Frequent predators | -0.01 ± 0.03 | - | - | - | - |
| Sex | 0.04 ± 0.02 | 1 | 37.52 | 5.45 | **0.03** |
| **(b) Daily growth of fish during the experiment** | | | | | |
| Intercept | 0.02 ± 0.01 | - | - | - | **-** |
| Feeding regime | -0.04 ± 5e^-3^ | 1 | 88.71 | 58.81 | **<0.01** |
| Predator exposure | - | 2 | 88.69 | 0.14 | 0.87 |
| Occasional predators | 3e^-3^ ± 0.01 | - | - | - | **-** |
| Frequent predators | 2e^-3^ ± 0.01 | - | - | - | **-** |
| Sex | 0.01 ± 5e^-3^ | 1 | 88.73 | 7.37 | **0.01** |

**Table S4:** (a) Daily weight gain and (b) daily growth of fish during the experiment. Fish were kept on two different feeding regimes (reduced or increased) and exposed to three frequencies of occurring predators (no predators, occasional predators, frequent predators). Measurements were done before the start of the feeding regimes and on the last day of each respective diet either just before the reversal learning task or the hormone sampling (depending on the fish). Results are shown from two linear mixed effect models. Fish on the increased diet gained more weight and grew more during the experiment than fish on the reduced diet whereas the frequency of predator exposures did not influence weight gain or growth of fish. Experimental treatments consisted of the independent manipulations of the subject’s nutritional state and predator experience. Diet was included as a two-level factor (increased, reduced) and the estimate is shown as difference to the reference level (‘increased’). Predator exposure was included as a three-level factor (‘no predator’, ‘occasional predators’, frequent predators’) and estimates are shown as difference to the reference level (‘no predators’). Sex was included as a two-level factor (‘male’, ‘female’) and the estimate is shown as difference to the reference level ‘female’. To obtain p-values a likelihood ration test was used to compare the models with and without the factor of interest. N=96 fish in 20 tanks and three blocks.

### Supplementary Table S5

| **Contrast** | **Estimate ± SE** | **z-value** | ***P*-value** |
| --- | --- | --- | --- |
| Intercept | 0.19 ± 0.57 | 0.35 | 0.73 |
| **(a) No predators vs. predators** | | | |
| (IN, RN) vs. (IO, IF, RO, RF) | -0.44 ± 0.29 | -1.52 | 0.13 |
| **(b) Frequent predators vs. occasional predators** | | | |
| (IO, IF) vs. (RO, RF) | 1.16 ± 0.53 | 2.19 | **0.03** |

**Table S5**: Probability of fish showing acute stress response to a novel stressor. Orthogonal contrasts are presented to investigate the significant main effect ‘Predator exposure’ in Table 1. Intercept estimate represents the grand mean of all treatments. First, we set the contrast of the model to compare the mean of all groups experiencing ‘no predators’ with the mean of all groups experiencing any predator exposure [(IN, RN) vs. (IF, IO, RF, RO)]. Second, we compared the mean of all groups experiencing ‘occasional predators’ with the mean of all groups experiencing ‘frequent predators’ [(IF, IO) vs. (RF, RO)]. Note that mean values of treatments presented in round brackets were used in the comparisons. Orthogonal comparisons of treatments (a, b) are displayed as: IN = increased – no predators; RN = reduced – no predators; IO = increased – occasional predators;. IF = increased – frequent predators; RO = reduced – occasional predators; RF = reduced – frequent predators. The direction of comparison within a contrast is left to right and the estimate value always refers to the treatment(s) to the right. If treatments are combined in parentheses, mean values of these treatments are used in the comparisons. N=34 fish from 20 groups and three blocks. p<0.05 is highlighted in bold.

### Supplementary Table S6

| **Factors** | **Estimate ± SE** | **Num. D.F.** | **Den. D.F** | **F-value** | ***P*-value** |
| --- | --- | --- | --- | --- | --- |
| Intercept | -28.27 ± 6.86 | - | - | - | - |
| Body condition | 9.31 ± 2.35 | 1 | 4.78 | 15.64 | **0.01** |

**Table S6**: The intensity of the stress reactivity after experiencing a novel stressor. Results are shown from a linear model with a gaussian error distribution. To obtain normally distributed residuals the dependent variable was log transformed. Fish body condition positively correlated with higher stress reactivity. For each fish cortisol concentrations were measured every 30 minutes (four samples) for two hours on the first water collection day (for more details see methods). The intensity of the stress reactivity was calculated as the difference between the sample showing peak cortisol concentrations and sample four (2h after novel stressor) and a positive value indicates a faster decrease in cortisol concentrations. Fish were exposed to frequent predator encounters and body condition was included as a continuous variable. To obtain p-values a likelihood ration test was used to compare the models with and without the factor of interest. N=34 fish from 20 tanks and 3 blocks. p<0.05 are highlighted in bold. N=11 fish from seven groups and three blocks. p<0.05 is highlighted in bold

### Supplementary Table S7

| **Factors** | **Estimate ± SE** | | **Χ^2^-value** | | ***P*-value** | |
| --- | --- | --- | --- | --- | --- | --- |
| Intercept | | 0.46 ± 0.7 | | - | | - |
| Body condition | | -0.25 ± 0.74 | | 0.19 | | 0.66 |
| Predator exposure | | -± | | 1.48 | | 0.48 |
| *Occasional predators* | | -0.4 ± 0.86 | | - | | - |
| *Frequent predators* | | 0.24 ± 0.93 | | - | | - |
| Sex | | 1.45 ± 0.83 | | 3.42 | | *0.06* |

**Table S7:** Probability of fish showing acute stress response on the fourth water sample collection day after fish were habituated to the new stressor. Results are shown from a generalized linear mixed effect model with a binomial error distribution. Body condition and predator exposure did not influence the probability to display an acute stress response. Fish were classified as acute or prolonged responders depending on their cortisol concentration profile for two hours on the fourth water collection day (see methods for more details). In short acute responders decreased cortisol concentrations within two hours whereas prolonged responders did not. Experimental treatments consisted of the independent manipulations of the subject’s nutritional state and predator exposure. Body condition was included as a two-level factor (high, low) and the estimate is shown as difference to the reference level (‘high’). Predator exposure was included as a three-level factor (‘no predator’, ‘occasional predators’, frequent predators’) and estimates are shown as difference to the reference level (‘no predators’). Sex was included as a two-level factor (‘male’, ‘female’) and the estimate is shown as difference to the reference level ‘female’. To obtain p-values a likelihood ration test was used to compare the models with and without the factor of interest. N=35 fish from 20 groups and three blocks.

### Supplementary Table S8

| **Contrast** | **Estimate ± SE** | **z-value** | ***P*-value** |
| --- | --- | --- | --- |
| Intercept | 3.42 ± 0.03 | 110.3 | **<0.01** |
| **(a) High body condition vs. low body condition** | | | |
| (IN, IO, IF) vs. (RN, RO, RF) | 0.04 ± 0.03 | 1.52 | 0.13 |
| **(b) No predators vs. predators (high body condition)** | | | |
| IN vs. (IO, IF) | -0.04 ± 0.03 | 1.32 | 0.19 |
| **(c) Occasional predators vs. frequent predators (high body condition)** | | | |
| IO vs. IF | -0.16 ± 0.05 | -3.4 | **<0.01** |
| **(d) No predators vs. predators (low body condition)** | | | |
| RN vs. (RO, RF) | -6e^-3^ ± 0.03 | -0.2 | 0.84 |
| **(e) Occasional predators vs. frequent predators (low body condition)** | | | |
| RO vs. RF | -6e^-3^ ± 0.06 | -0.11 | 0.91 |

**Table S8:** The number of trials until fish reached the learning criterion in the reversal phase of the learning task. Orthogonal contrasts are presented which were done after confirming that the interaction ‘Body condition x Predator exposure’ was p<0.1 in Table 3. Intercept estimate represents the grand mean of all treatments. First, we set the contrast of the model to compare the mean of all fish with a high body condition with the mean of all fish with a low condition [(IN, IO, IF) vs. (RN, RO, RF)]. Second, we only used fish with a high body condition to compare fish that experienced no predators with the mean of fish that experienced any predator exposure [IN vs. (IO, IF)]. Third, we only used fish with a high body condition to compare fish that experienced occasional predators with fish that experienced frequent predators [IO vs. IF]. Fourth, we only used fish with a low body condition to compare fish that experienced no predators with the mean of fish that experienced any predator exposure [RN vs. (RO, RF)]. Fifth, we only used fish in a low body condition to compare fish that experienced occasional predators with fish that experienced frequent predators [RO vs. RF]. Note that mean values of treatments presented in round brackets were used in the comparisons. Orthogonal comparisons of treatments (a-e) are displayed as: IN = increased – no predators; IO = increased – occasional predators; IF = increased – frequent predators; RN = reduced – no predators;. RO = reduced – occasional predators; RF = reduced – frequent predators. The direction of comparison within a contrast is left to right and the estimate value always refers to the treatment(s) to the right. If treatments are combined in parentheses, mean values of these treatments are used in the comparisons. N=34 fish from 20 groups and three blocks. p<0.05 are highlighted in bold.

### Supplementary Table S9

| **Factors** | **Estimate ± SE** | **Χ^2^-value** | ***P*-value** |
| --- | --- | --- | --- |
| (**a) Error rates in the acquisition phase** | | | |
| Intercept | -1.37 ± 0.26 | - | - |
| Body condition | 0.13 ± 0.15 | 0.73 | 0.39 |
| Predator exposure | - | 3.4 | 0.18 |
| *Occasional predators* | 0.34 ± 0.24 | - | - |
| *Frequent predators* | 0.55 ± 0.34 | - | - |
| **(b) Error rates in the reversal phase** | | | |
| Intercept | -0.85 ± 0.15 | - | - |
| Body condition | -0.06 ± 0.11 | 0.27 | 0.6 |
| Predator exposure | - | 0.24 | 0.89 |
| *Occasional predators* | -0.05 ± 0.15 | - | - |
| *Frequent predators* | -0.08 ± 0.17 | - | - |

**Table S9:** Error rates of fish during the (a) acquisition phase and (b) reversal phase of the learning task. Results are shown from two generalized linear mixed effect model with binomial error distributions. Error rates were calculated as the overall number of correct and wrong trials and included as a two-vector proportional response variable. Body condition and predator exposure did not influence the error rates of fish during the acquisition or reversal phase. Experimental treatments consisted of the independent manipulations of the subject’s nutritional state and predator exposure. Body condition was included as a two-level factor (high, low) and the estimate is shown as difference to the reference level (‘high’). Predator exposure was included as a three-level factor (‘no predator’, ‘occasional predators’, frequent predators’) and estimates in are shown as difference to the reference level (‘no predators’). To obtain p-values a likelihood ration test was used to compare the models with and without the factor of interest. (a) N=55 fish from 20 groups and three blocks; (b) N=56 fish from 20 groups and three blocks.

### Supplementary Figure S1

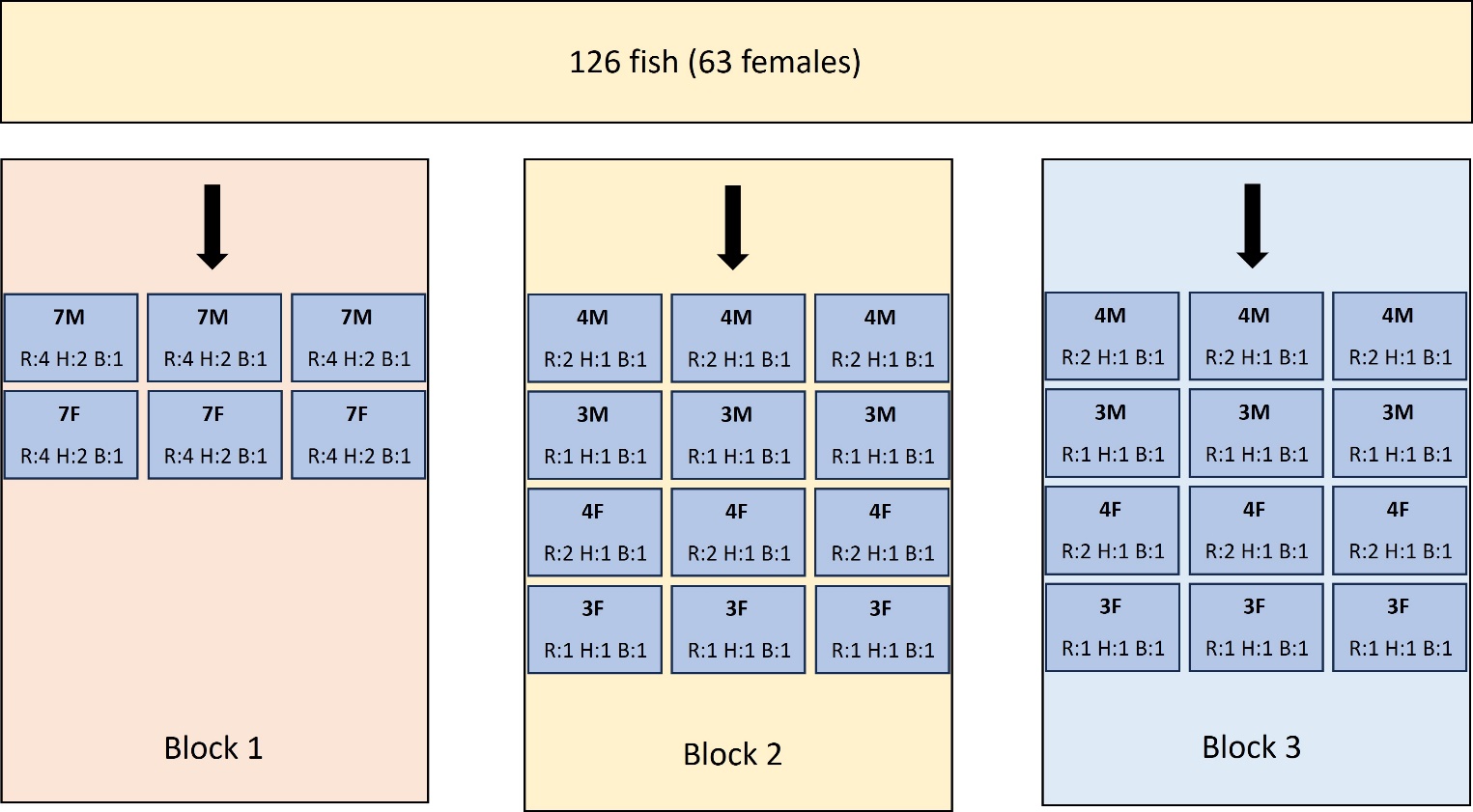

**Fig. S1:** Overview of experimental groups. Overall, we used 126 fish (63 females) in three blocks that were separately run. In each block, fish were kept in tanks with group sizes of seven (block 1), four (block 2, 3) or three (block 2, 3). Each group consisted of individuals of the same sex (M= male, F=female). From each group we randomly chose fish for the reversal learning task (R), Hormone sampling (H), or brain collection (B). Please note that fish used for brain collection are not part of this study and results will be reported elsewhere.

### Supplementary Figure S2

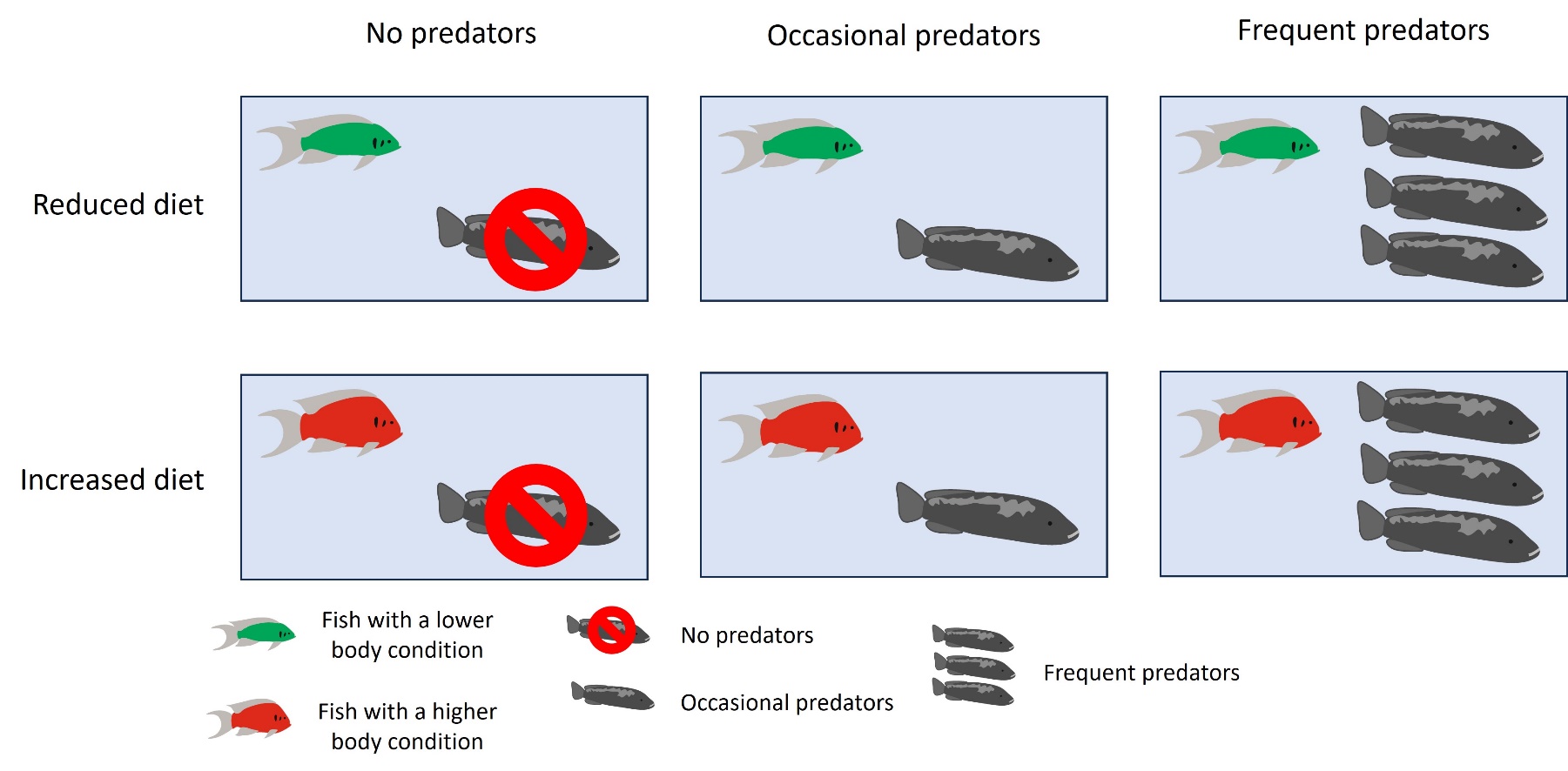

**Fig. S2:** Overview of treatments. To manipulate the nutritional state and the frequency of exposure to a stressor we kept fish either on a reduced diet (upper row) or an increased diet (lower row) and three different predator exposures: a control condition without any predator exposure (left column), an occasional exposure (middle column) or a frequent exposure (right column). To standardize predator presentations, we used video playbacks of attacking predators together with standardized stress manipulation and the transfer of olfactory cues (see methods for more details).

### Supplementary Figure S3

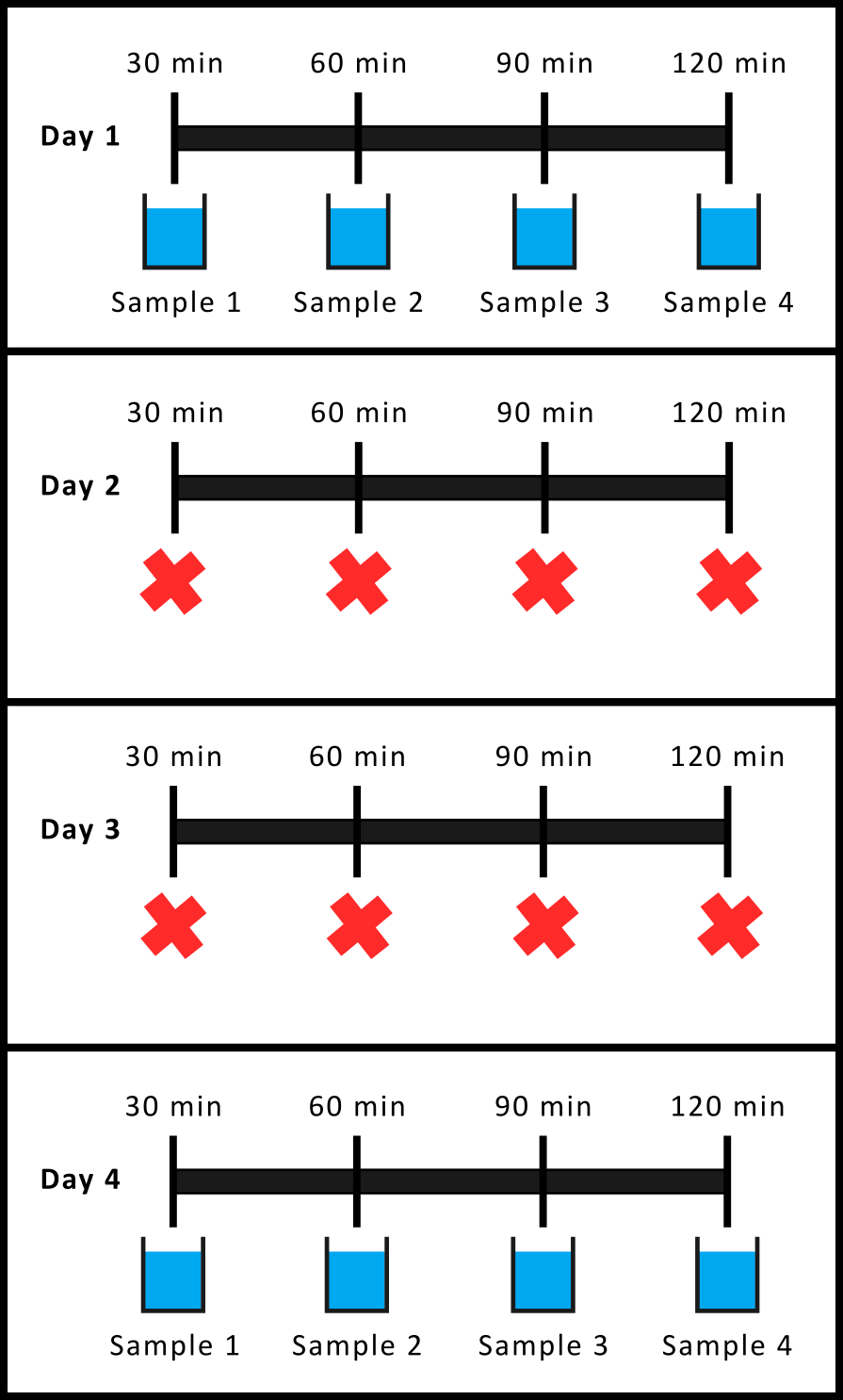

**Fig. S3**: Timeline of the hormonal sampling procedure. On day one, two fish were caught from the group tanks and transferred to individual sampling containers. For the next two hours four samples were collected using the flow-through system (more details in Methods). After the last water sample was collected (sample 4) both fish were returned to their respective group tanks. In the following two days we repeated the same procedure but did not collect any water samples. This ensured that test fish habituated to the procedures and allowed us to collect water samples of habituated fish. On the fourth day, the same procedures as on day 1 were repeated and we collected another four water samples per fish.

### Supplementary Figure S4

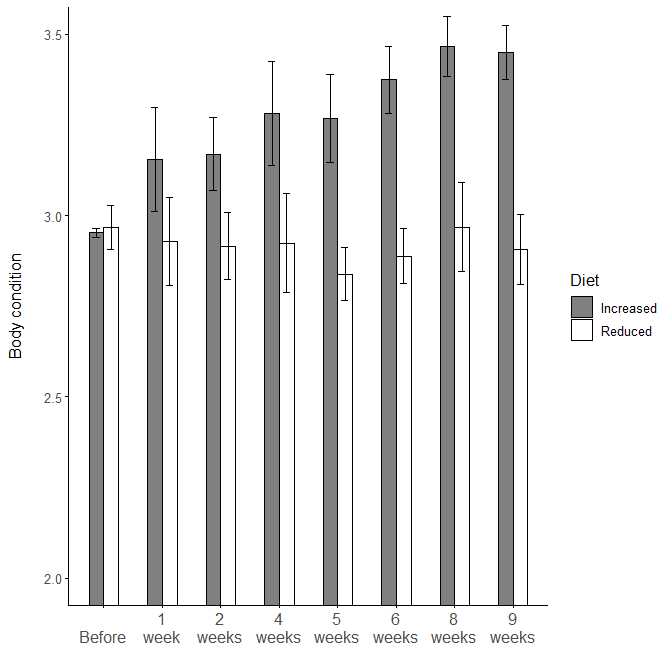

**Fig. S4**: Average (± SE) body conditions of pilot fish on two different diets (increased, reduced) during the first phase of the pilot experiment. Dark bars represent data from five fish on the increased diet (1.5 sticks per fish of JBL Stick M for cichlids©) and white bars represent data from four fish on the reduced diet (0.5 sticks of JBL Stick M for cichlids©). Body conditions were not different between the groups before the feeding regime started but quickly diverged afterwards. Fish were fed on their respective diets for nine weeks and throughout the first phase of the pilot experiment fish on an increased diet had higher body conditions than fish on a reduced diet. See Table S1 for statistical analysis

### Supplementary Figure S5

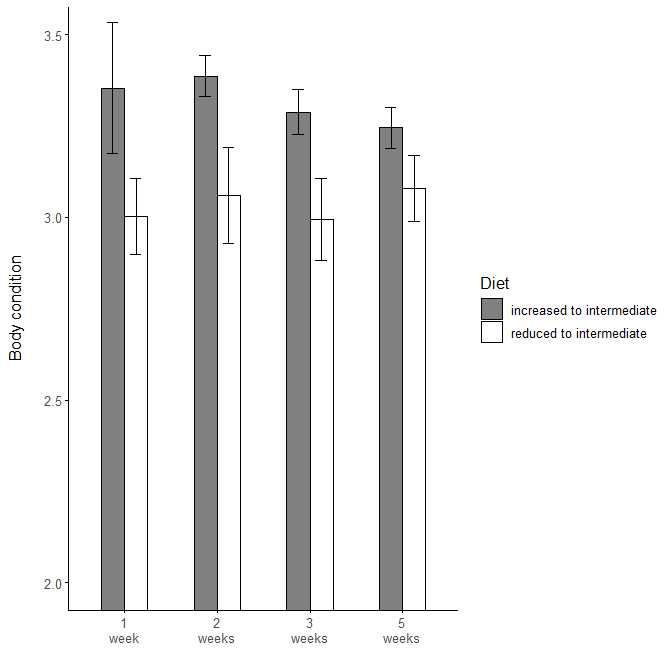

**Fig. S5:** Average (± SE) body condition of pilot fish after the switch to an intermediate (1 stick per fich JBL Stick M for cichlids©) diet in the second phase of the experiment. Dark bars represent data from five fish that were previously on an increased diet and white bars represent data from four fish that were previously on a reduced diet. Fish were fed the intermediate diet for five weeks and throughout the entire second phase of the experiment fish that were previously on an increased diet had higher body condition than fish previously on a reduced diet. See Table S2 for statistical analysis.

### Supplementary Figure S6

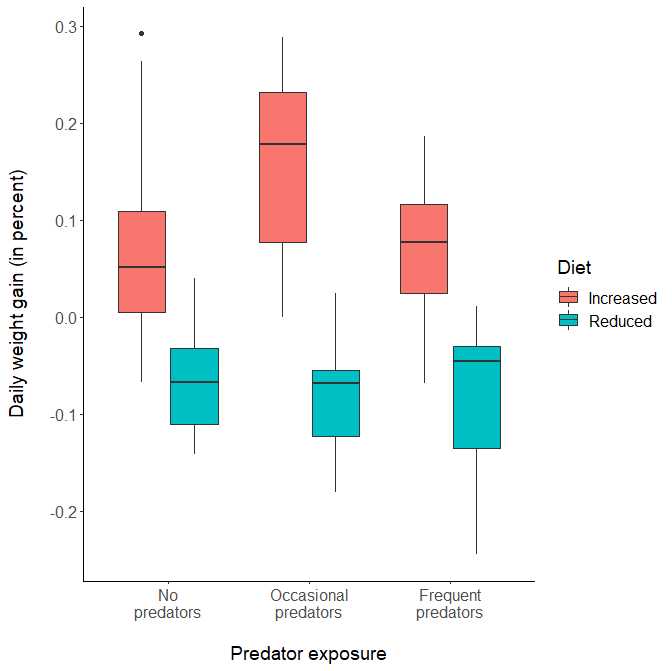

**Fig. S6:** Daily weight gain (in percent) of fish during the experiment. Fish were on two different diets (increased [red boxes] or reduced [blue boxes]) and experienced three types of predator exposures (no predators, occasional predators or frequent predators). Fish were measured before the start of the feeding regime and at the last day of the respective diets just before the reversal learning task or hormone sampling. Fish on the increased diet gained more weight during the course of experiment than fish on the reduced diet. The frequency of the predator exposure did not influence the daily weight gain. Shown are boxplots where boxes represent all data points within the lower .25 and higher .75 quartile. Thick horizontal lines represent the median, whiskers include all data points within 1.5*the length of the box (interquartile range) and dots represent outliers. See Table S4a for statistical analysis.

### Supplementary Figure S7

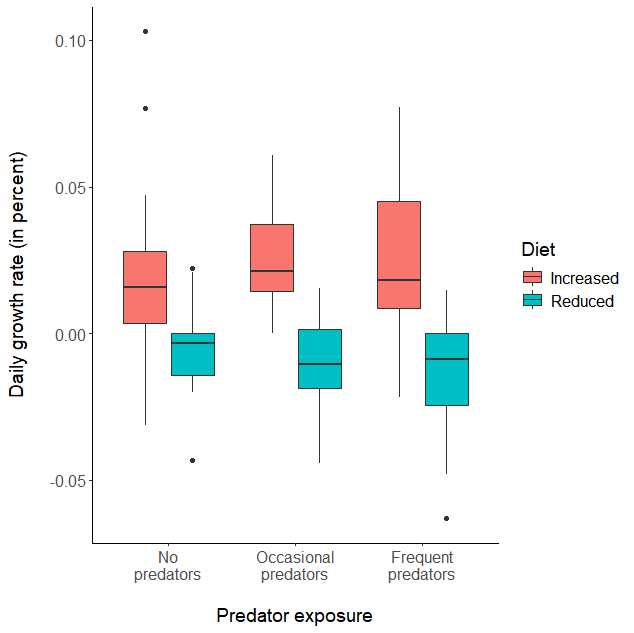

**Fig. S7**: Daily growth rate (in percent) of fish during the experiment. Fish were on two different diets (increased [red boxes] or reduced [blue boxes]) and experienced three types of predator exposures (no predators, occasional predators or frequent predators). Fish were measured before the start of the feeding regime and at the last day of the respective diets just before the reversal learning task or hormone sampling. Fish on the increased diet grew more during the course of experiment than fish on the reduced diet. The frequency of the predator exposure did not influence the daily growth rate. Shown are boxplots where boxes represent all data points within the lower .25 and higher .75 quartile. Thick horizontal lines represent the median, whiskers include all data points within 1.5*the length of the box (interquartile range) and dots represent outliers. See Table S4b for statistical analysis.

### Supplementary Figure S8

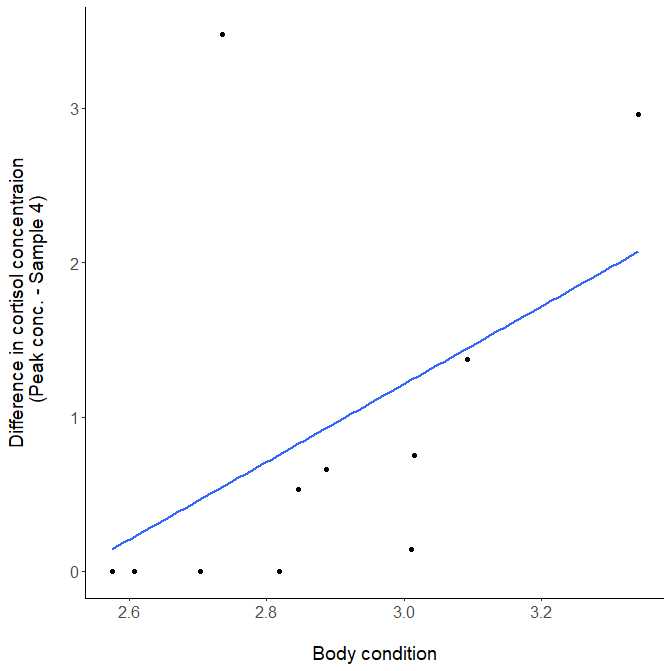

**Figure S8:** The ability to terminate the stress response of fish that were exposed to frequent predators (N=11). A stress recovery index was calculated as the difference between the peak cortisol concentration and the concentration at sample point four (2h after the novel stressor). A higher index indicates a rapid decrease of cortisol levels within two hours after a novel stressor and thus, a strong resilience. Zero values indicate that the peak concentration and the concentration at sample point four were equal, i. e. no decrease in the cortisol pattern within two hours. Body condition was positively correlated with the stress reactivity index. For statistical analysis see Table 2.

### Supplementary references

[1] Jungwirth, A., Balzarini, V., Zottl, M., Salzmann, A., Taborsky, M. & Frommen, J.G. 2019 Long-term individual marking of small freshwater fish: the utility of Visual Implant Elastomer tags. *Behav. Ecol. Sociobiol.* **73**. (doi: 10.1007/s00265-019-2659-y).

[2] Scott, A.P. & Ellis, T. 2007 Measurement of fish steroids in water--a review. *Gen Comp Endocrinol* **153**, 392-400. (doi: 10.1016/j.ygcen.2006.11.006).

[3] Scott, A.P., Hirschenhauser, K., Bender, N., Oliveira, R., Earley, R.L., Sebire, M., Ellis, T., Pavlidis, M., Hubbard, P.C., Huertas, M., et al. 2008 Non-invasive measurement of steroids in fish-holding water: important considerations when applying the procedure to behaviour studies. *Behaviour* **145**, 1307-1328. (doi: 10.1163/156853908785765854).

[4] Ellis, T., James, J.D., Stewart, C. & Scott, A.P. 2004 A non-invasive stress assay based upon measurement of free cortisol released into the water by rainbow trout. *J. Fish Biol.* **65**, 1233-1252. (doi: 10.1111/j.0022-1112.2004.00499.x).
